# The rediscovery of a relict unlocks the first global phylogeny of whip spiders (Amblypygi)

**DOI:** 10.1101/2022.04.26.489547

**Authors:** Gustavo S. de Miranda, Siddharth S. Kulkarni, Jéssica Tagliatela, Caitlin M. Baker, Alessandro P.L. Giupponi, Facundo M. Labarque, Efrat Gavish-Regev, Michael G. Rix, Leonardo S. Carvalho, Lívia Maria Fusari, Hannah M. Wood, Prashant P. Sharma

**Author notes:** Equal author contribution.

## Abstract

Asymmetrical rates of cladogenesis and extinction abound in the Tree of Life, resulting in numerous minute clades that are dwarfed by larger sister groups. Such taxa are commonly regarded as phylogenetic relicts or “living fossils” when they exhibit an ancient first appearance in the fossil record and prolonged external morphological stasis, particularly in comparison to their more diversified sister groups. Due to their special status, various phylogenetic relicts tend to be well-studied and prioritized for conservation. A notable exception to this trend is found within Amblypygi (“whip spiders”), a visually striking order of functionally hexapodous arachnids that are notable for their antenniform first walking leg pair (the eponymous “whips”). Paleoamblypygi, the putative sister group to the remaining Amblypygi, is known from Late Carboniferous and Eocene deposits, but is survived by a single living species, *Paracharon caecus* Hansen, 1921, that was last collected in 1899. Due to the absence of genomic sequence-grade tissue for this vital taxon, there is no global molecular phylogeny for Amblypygi to date, nor a fossil-calibrated estimation of divergences within the group. Here, we report several individuals of a previously unknown species of Paleoamblypygi from a cave site in Colombia. Capitalizing upon this discovery, we generated the first molecular phylogeny of Amblypygi, integrating ultraconserved element sequencing with legacy Sanger datasets and including described extant genera. To quantify the impact of sampling Paleoamblypygi on divergence time estimation, we performed *in silico* experiments with pruning of *Paracharon*. We demonstrate that the omission of relicts has a significant impact on the accuracy of node dating approaches that outweighs the impact of excluding ingroup fossils. Our results underscore the imperative for biodiversity discovery efforts in elucidating the phylogenetic relationships of “dark taxa”, and especially phylogenetic relicts in tropical and subtropical habitats.

## Introduction

A phylogenetic relict represents the remnant of a previously more diverse fauna that has undergone faunal turnover and extinction events (Simpson 1955; Darlington 1957; Holmquist 1962; Gould et al. 1977). The evidence for this condition often consists of an ancient appearance of a lineage in the fossil record and an historically broad distribution of fossils, with the retention of only a small number of geographically-restricted extant species (Grandcolas et al. 2014).

Typically, phylogenetic relicts represent an example of extreme phylogenetic attenuation, either as a result of extinction, low speciation rate, or a combination of these two processes. Relicts are often the sister taxa of diverse clades, which confers a special status upon these small groups for evolutionary and genomic studies, as well as conservation priority. Renowned examples of phylogenetic relicts include coelocanths, lungfishes, tuataras, monotremes, nautiloids, horseshoe crabs, gingko, and *Amborella* trees (Grandcolas et al. 2014; 2016).

Recapitulating a historical fascination with the causal mechanisms that create relicts, modern investigations of phylogenetic relicts have focused on evidence of bradytelic evolution at the level of genes and genome architecture, as well as the signatures of extinction and faunal turnover (Soltis et al. 2008; Amemiya et al. 2010, 2013; Combosch et al. 2017; Meyer et al. 2021; Nong et al. 2021). Regardless of the cause of their taxonomic attenuation, the significance of phylogenetic relicts is twofold from the perspective of phylogenetics. First, as the established sister groups of more diverse clades, such relicts play an outsized role in polarizing character states and reconstructing patterns of morphological evolution and genome organization. Second, as an extension of both topological placement and their fossil record, phylogenetic relicts play vital roles in molecular dating, specifically through the provision of calibration points for node dating.

Given the attention and research focus that many phylogenetic relicts command, obtaining genomic and phylogenetic representation for such groups is often prioritized, even in cases where these taxa are geographically restricted (e.g., *Amborella*; tuataras; *Cryptocercus* cockroaches; Inward et al. 2007; Soltis et al. 2008; Gemell et al. 2020). One notable exception to this trend is the monotypic family Paracharontidae, the sister group to the remaining Amblypygi, commonly known as whip spiders. The order Amblypygi is comprised of ca. 260 species, yet has a broad global distribution across tropical and subtropical habitats (Weygoldt, 1996; Harvey 2003; Miranda et al. 2022). Whip spiders are notable for the fearsome appearance of their hypertrophied raptorial pedipalps, as well as the modification of the first walking leg pair into antennal analogs—these arachnids are functionally hexapodous (Fig. 1). Amblypygi have also featured prominently in studies of arachnid behavior and communication, as they exhibit complex behaviors such as homing, learning, and aggregation as a function of kinship groups (Fowler-Finn and Hebets 2006; Wiegmann et al. 2016; Flanigan et al. 2021).

**Figure 1.**
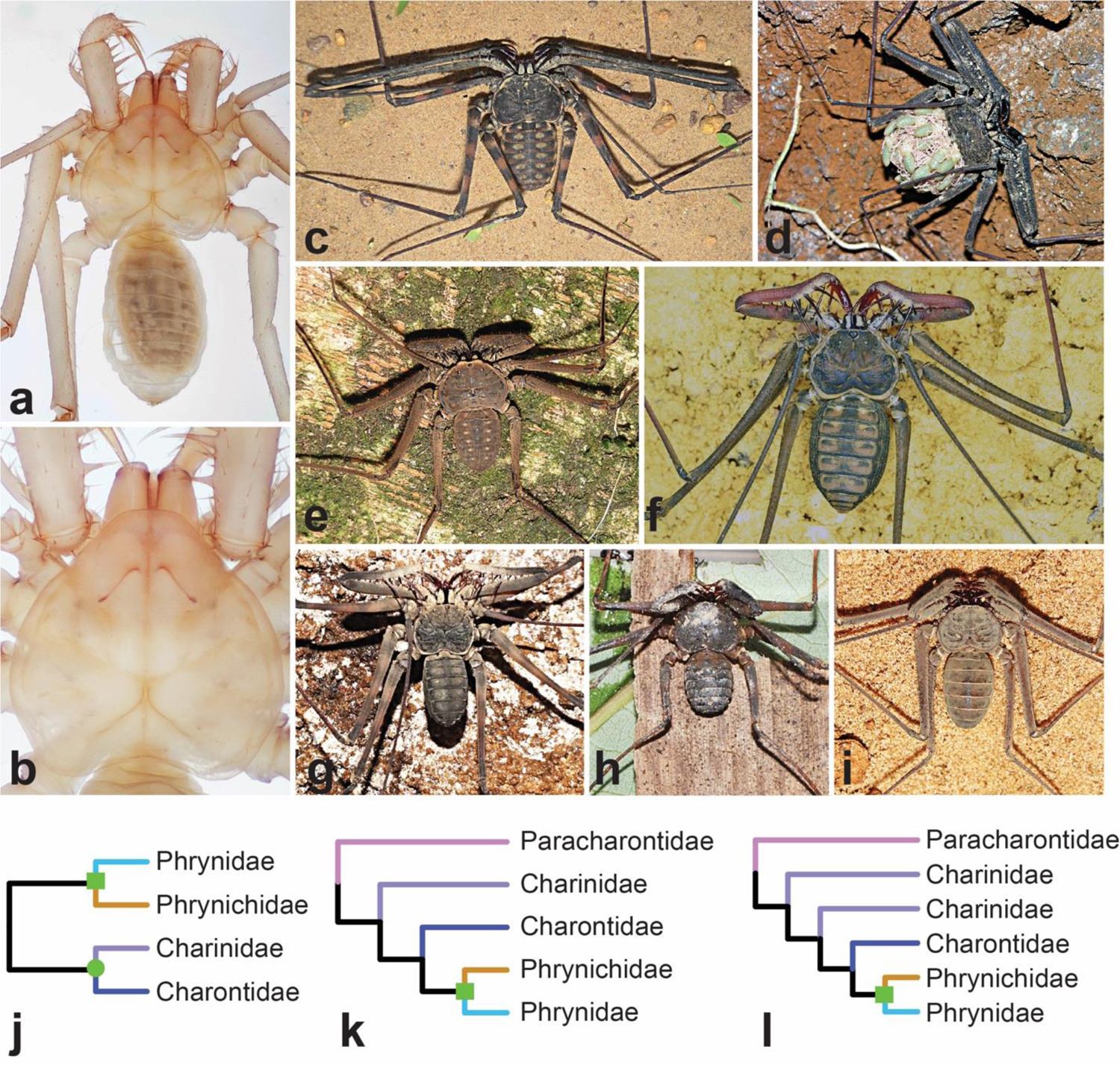
Exemplars of whip spider diversity and overview of morphology-based classification schema. a) Habitus of adult *Paracharon* sp. from Colombia. Note orthognathous articulation of the pedipalps. b) Magnification of the prosoma of *Paracharon* sp., highlighting the absence of eyes. c*) Trichodamon princeps*. d) Adult female of *Damon medius*, carrying juveniles on the dorsal opisthosoma. e) *Paraphrynus aztecus*. f) *Heterophrynus batesii*. g) *Catageus* sp. h) *Sarax* sp. i). *Charinus cearensis* j) Cladogram from Quintero (1986). Genera have been replaced with current familial assignments. k) Interfamilial relationships inferred by Weygoldt (1996). l) Interfamilial relationships inferred by Garwood et al. (2017). Green squares and circle represent.

Whereas higher-order (and often, genome-scale) molecular phylogenies are now available for almost all extant chelicerate orders (Klompen et al. 2007; Giribet et al. 2014; Clouse et al. 2017; Fernández et al. 2017; Wheeler et al. 2017; Klimov et al. 2018; Benavides et al. 2019, 2021; Ballesteros et al. 2021; Santibáñez-López et al. 2022), a global molecular phylogeny for Amblypygi remains unavailable (Miranda et al. 2022). This gap is specifically attributable to the elusive nature of the putative sister group of the remaining extant whip spiders; the family Paracharontidae was known from a single species, *Paracharon caecus*, described one century ago from two localities in modern day Guinea Bissau (Hansen 1921). Morphological phylogenetic analyses have suggested that the oldest known crown group whip spider fossils, from Carboniferous deposits in North America and the British Isles, constitute stem groups of Paracharontidae (Garwood et al. 2017). A more recent discovery of this lineage in Cambay Basin amber in South Asia supports a historically broader distribution of Paracharontidae across the Paleotropics by the Eocene (Engel and Grimaldi 2014). Fossils of the remaining extant Amblypygi taxa (Euamblypygi) are comparatively recent, dating between the Cretaceous and the Miocene (Engel and Grimaldi 2014).

Such patterns of early branching phylogenetic relicts are common within Chelicerata (Supplementary Table S1). As examples, mesothele spiders (ca. 0.36% of described spider diversity) once had a broad geographic range including Western Europe in the Carboniferous, but are presently restricted to parts of east and southeast Asia (Selden 1996, Kallal et al. 2021). Cyphophthalmi, the cryptic sister group of the remaining Opiliones (ca. 3.1% of described harvestman diversity), exhibit distributions, internal relationships, and molecular divergence times that nearly mirror the sequence and timing of supercontinental fragmentation (Boyer et al. 2007; Giribet et al. 2012; Baker et al. 2020). But in the case of Amblypygi, the absence of the ‘living fossil’ *Paracharon* in any recent historical collections has effectively barred any higher-level molecular phylogenetic assessment of this group, as well as molecular dating that leverages the availability of fossil data. A handful of morphological cladistic analyses have tackled whip spider relationships, but these exhibit marked incongruence in higher-level nodes (Quintero 1986; Weygoldt 1996; Garwood et al. 2017; Fig. 1). This incongruence is partly attributable to the morphological conservatism of extant whip spiders, which incurs a paucity of discrete variable characters in the group.

Propitiously, we discovered an undescribed species of *Paracharon* from a collection at a cave in Colombia, consisting of one adult and two juvenile specimens. The rediscovery of this lineage after 123 years opens the door to the first higher-level molecular phylogeny of this poorly understood arachnid group. Here, we present a global molecular phylogeny of Amblypygi using ultraconserved element sequencing in tandem with Sanger datasets, and sampling all extant genera. To quantify the impact of this relict lineage on evolutionary analyses, we performed multiple divergence time estimations with and without *Paracharon* sequence data, as well as with and without Amblypygi fossils. We show that the sampling of *Paracharon* plays a more significant role in the accuracy of divergence time estimation than the use of ingroup calibration fossils. Our results underscore the significance of phylogenetic relicts in evolutionary analyses.

## Materials and Methods

### Species sampling

Specimens sequenced for this study were hand collected from field sites or contributed by collections of the National Museum of Natural History, the Smithsonian Institution, Washington, DC, USA (USNM); the Museum of Comparative Zoology, USA (MCZ); the Natural History Museum of Denmark, Copenhagen (NHMD); the Museu Nacional do Rio de Janeiro, Brazil (MNRJ); the California Academy of Sciences, San Francisco, USA (CAS); the State Museum of Natural History Stuttgart, Germany (SNMS); and the Universidade Federal do Piauí, Brazil (CHNUFPI). We selected exemplars of each of the five described extant families, and all 17 described genera: 20 specimens of Charinidae (133 described species), one Charontidae (15 described species), 1 Paracharontidae (1 described species), two Phrynichidae (35 described species), and ten Phrynidae (77 described species). Outgroup sampling leveraged previous phylogenomic works that have established Amblypygi as the sister group of vinegaroons (Thelyphonida), and short-tailed whip scorpions (Schizomida) (the clade Pedipalpi; Sharma et al. 2014; Ballesteros and Sharma 2019; Ballesteros et al. 2019). Pedipalpi in turn is understood to form a clade with spiders, scorpions, and pseudoscorpions, a relationship supported by rare genomic changes (e.g., Ontano et al. 2021, 2022; Ballesteros et al. 2022). We therefore included two representatives of spiders, one vinegaroon, three Schizomida, and rooted the tree with three scorpions. Accession data for all specimens in this study are provided in Supplementary Table S2.

### Ultraconserved element sequencing

Ultraconserved element (UCE) sequences generated in this study were augmented with published UCE and RNASeq datasets (Supplementary Table S3). We analyzed 47 terminals with UCE data including 38 Amblypygi. For newly sequenced specimens, one leg was used for DNA extractions from one specimen using the DNeasy™ Tissue Kit (Qiagen Inc., Valencia, CA). Libraries were prepared and enriched following protocols in Faircloth et al. (2015), but following the modifications detailed in the Supplementary Material. All pools were enriched with the Arachnida probe set (Starrett et al. 2017) except *Heterophrynus* and *Charon*, which were enriched using the Spider2Kv1 probe set (Kulkarni et al. 2020) following the myBaits protocol 4.01 (Arbor Biosciences).

### Inclusion of Sanger-sequenced terminals

While UCE datasets have demonstrated great potential for leveraging historical collections, the compatibility of these data with legacy datasets in Sanger-sequencing studies is often not evaluated. To integrate historical datasets in a comprehensive phylogenetic framework, we compiled another matrix comprised of 109 terminals for six publicly available Sanger-sequenced loci: 12S ribosomal RNA (12S), 16S rRNA (16S), cytochrome *c* oxidase subunit 1 (COI), histone H3 (H3), and the small and large subunits of nuclear ribosomal genes (18S and 28S, respectively). We integrated this data set by querying the raw files of the UCE data set for any potential match with 18S rRNA and 28S rRNA. Finally, COI and H3 markers were aligned with peptide translation using MACSE (Ranwez et al. 2011), with the invertebrate mitochondrial code implemented for COI. The remaining markers (12S, 16S, 18S and 28S) were aligned using MAFFT version 7 (Katoh and Standley 2013). Trimming was performed on all alignments using trimAL (Capella-Gutiérrez et al. 2009) with *-gappyout*.

### Phylogenomic analyses

The assembly, alignment, trimming and concatenation of data were done using the PHYLUCE pipeline (publicly available at https://phyluce.readthedocs.io/en/latest/). UCE contigs were extracted using the Spider2Kv1 probe set (Kulkarni et al. 2020) and the Blended probe set (Maddison et al. 2020). We applied gene occupancies of 10%, 25% and 40% on the UCE data set. Additionally, we also analyzed 1% occupancy of the UCE data set to allow inclusion of all loci in the reconstruction of the phylogeny. We screened for orthologous and duplicate loci with the minimum identity and coverage of 65 and 65 matches.

To augment the UCE dataset with RNASeq datasets, we followed the assembly, sanitation and reading frame detection pipeline as in Fernández et al. (2018) for assembling the transcriptomes. Additionally, we ran the Perl script for Rcorrector (Song and Florea, 2015) for error correction and downstream efficiency prior to assembly. FASTA files of transcriptomes resulting from CD-HIT-EST were converted to 2bit format using faToTwoBit, (Kent et al. 2002). In the PHYLUCE environment (publicly available at https://phyluce.readthedocs.io/en/latest/tutorial-three.html), we created a temporary relational database to summarize probe to assembly match using: *phyluce_probe_run_multiple_lastzs_sqlite* function on the 2bit files. The ultraconserved loci were recovered by the *phyluce_probe_slice_sequence_from_genomes* command. The resulting FASTA files were treated as contigs and used to match the reads to the Blended probe set of Maddison et al. (2020).

Phylogenetic analyses were performed on two types of data sets: an unpartitioned UCE nucleotide data set alone (at difference thresholds of occupancy); and a matrix of unpartitioned UCE paired with the partitioned six Sanger loci matrix. Maximum likelihood analyses were performed using IQ-TREE (Nguyen et al. 2015) version 2. Model selection was allowed for each unpartitioned dataset using the TEST function (Kalyaanamoorthy et al. 2018, Hoang et al. 2018). Nodal support was estimated via 1,000 ultrafast bootstrap (UFBoot) replicates (Hoang et al. 2018) and Shimodaira-Hasegawa approximate likelihood ratio testing (SH-aLRT) with 1,000 iterations. We used the flag *-bnni* which reduces the risk of overestimating branch support with UFBoot due to model violations. This flag optimizes each bootstrap tree using a hill-climbing nearest neighbor interchange (NNI) search based on the corresponding bootstrap alignment (Hoang et al. 2018). We used gene (gCF) and site concordance factors (sCF) to evaluate the percentage of gene trees and decisive alignment sites containing a given branch in the maximum likelihood tree implemented in IQ-TREE (Minh et al. 2020).

To evaluate topological robustness of selected nodes, we evaluated signal for alternative placements using quartet likelihood mapping (Strimmer and Von Haeseler 1997) in IQ-TREE. Likelihood mapping was performed against the complete dataset, representing all UCE loci (i.e., not filtered for any occupancy threshold).

### Phylogenomic dating

Divergence time estimation was performed using a Bayesian inference approach, as implemented in codeml and MCMCTree (both part of PAML v.4.8; Yang 2007; dos Reis and Yang 2019). We optimized the fossil information-based calibrations on a matrix comprising smallest UCE matrix (selected for highest taxon occupancy), plus the six Sanger loci. The maximum likelihood tree topology inferred for this dataset served as the basis for divergence time estimation, implementing uniform node age priors to accommodate the scarcity of terrestrial chelicerate fossils. The root age was set to a range of 545 Mya to 435 Mya, based on the age of *Eramoscorpius brucensis*, the oldest known arachnopulmonate. *Weygoldtina anglica*, the oldest known stem-Palaeoamblypygi, was used to constrain the most recent common ancestor (MRCA) of crown Amblypygi. The Cretaceous fossils *Kronocharon prendinii* and *Britopygus weygoldtii* were assigned to the MRCA of (Charinidae + Charontidae) and (Phrynidae + Phrynichidae), respectively. Four other outgroup nodes were calibrated using the oldest unambiguous fossils representing those clades. Justifications and references for node calibrations are provided in Supplementary Table S3. Both chains were run for 20,000 generations, with an additional 5% set for burnin. We used the independent rates clock model for all partitions and all analyses were run using the same *seed* value (arbitrarily set to 5) to make the results reproducible.

Separately, we inferred divergence times under a penalized likelihood approach, as implemented in LSD2, which uses a least-squares approach based on a Gaussian model and is robust to uncorrelated violations of the molecular clock (To et al. 2016). The root age was set to a maximum of 545 Ma (Supplementary Table S3). We used the commands *prime* and *thorough* to optimize the analyses, and cross validation was used to select the optimal smoothing parameter. Following Eberle et al. (2018), penalized likelihood optimization iterations were increased from the default of 2 to 5, and the number of penalized likelihood simulated annealing was doubled from 5,000 to 10,000. LSD2 requires at least one fixed date, so we used 314.6 Mya as a fixed date for the most recent common ancestor of Scorpiones.

To investigate the influence of the relict *Paracharon* on divergence time estimation (i.e., simulate the effect of not having discovered the Colombian specimens), we performed a separate family of analyses for both dating approaches (MCMCTree and LSD2), wherein we removed the terminal *Paracharon* sp.; the MRCA of the remaining Amblypygi (Euamblypygi) was therefore calibrated with the minimum age constraint reflecting the age of the Carboniferous fossil *Weygoldtina anglica.* We compared the median age and variance of nodes between runs to assess which parts of the chronogram were most prone to node age overestimation. In addition, to compare the significance of sampling *Paracharon* versus ingroup fossil calibrations, we performed a third family of analyses wherein we retained all terminals, but removed the three ingroup fossil node calibrations and keeping only the five calibration points for outgroups.

## Results

### Phylogenomics

We reconstructed an Amblypygi phylogeny using UCE datasets, as well as a combination of UCEs and six Sanger loci. Phylogenetic relationships obtained from UCE datasets with varying levels of missing data (10%, 25%, and 40% occupancy) were grossly similar (Fig. 2; Supplementary Fig. S1). The 40% occupancy data set contained 135 loci and represented the least amount of missing data, and was therefore combined with the six-locus Sanger dataset for dating analyses. Six-locus alignments were combined with 44 out of 48 UCE alignments (Supplementary Table S4).

**Figure 2.**
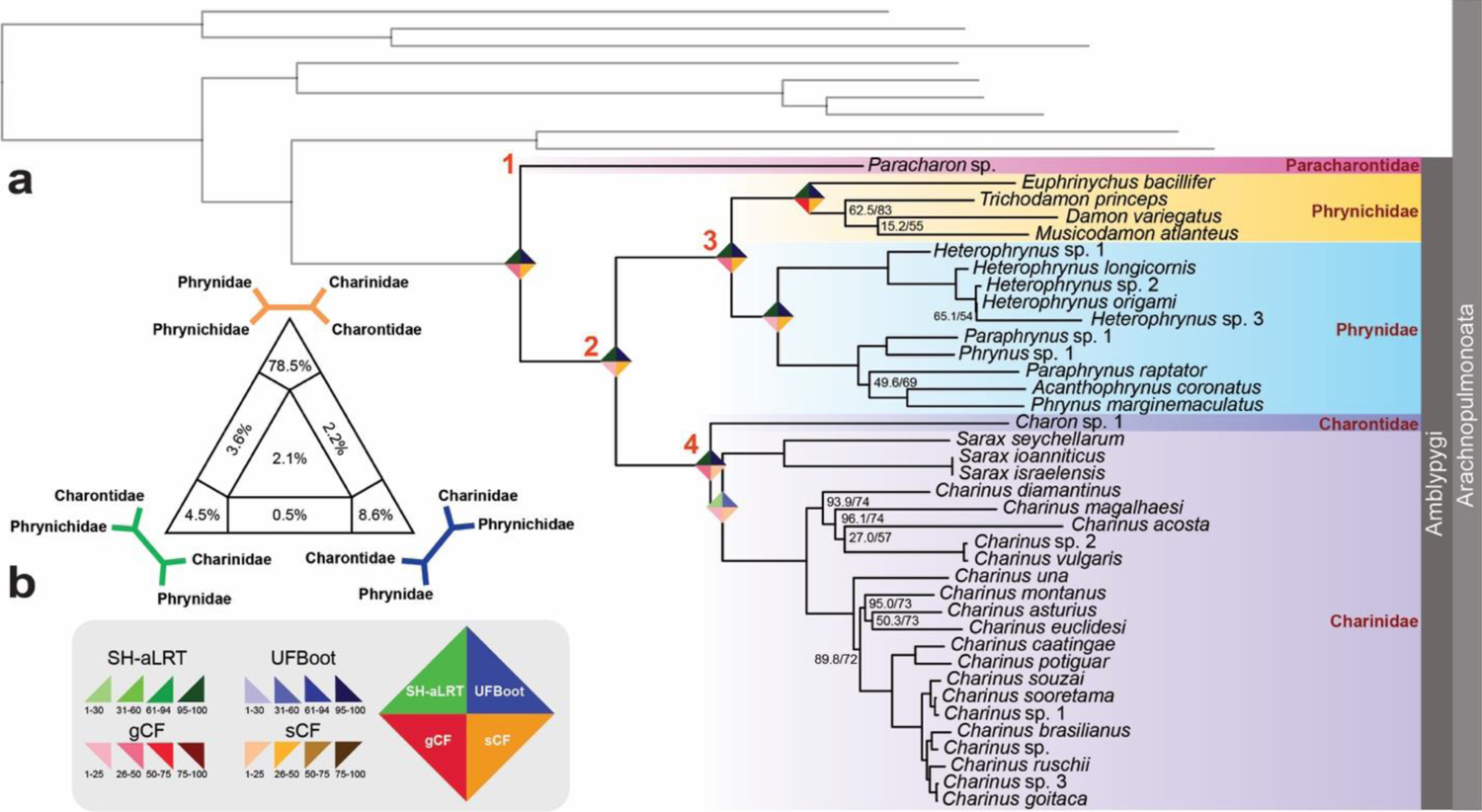
a) Phylogenomic relationships of Amblypygi based on 135 UCE loci (40% occupancy). Numbers on nodes correspond to SH-aLRT / UFBoot, respectively; unmarked nodes are maximally supported. Rhombus icons on selected nodes reflect nodal support values, corresponding to the legend in (b). b) Four-cluster likelihood mapping assessing support for Charontida (Charontidae + Charinidae), based on the supermatrix of all UCE loci. Red numbers indicate higher-level taxa: 1, Paleoamblypygi; 2, Euamblypygi; 3, Phrynoidea; 4, Charontoidea.

All datasets recovered Paracharontidae as sister to all other extant Amblypygi (Euamblypygi), as proposed by Weygoldt (1996) (Figs. 2, 3). Together with some extinct taxa, Paracharontidae forms Paleoamblypygi (Weygoldt, 2000; Garwood et al., 2017), distinguished from the remaining whip spiders (i.e., Euamblypygi) by the dorso-ventral articulation of the pedipalps and the anterior projection of the carapace (Weygoldt, 1996; Garwood et al. 2017). Phrynichidae and Phrynidae (Phrynoidea) were recovered in a clade sister to Charinidae plus Charontidae (Charontoidea). Phrynoidea and Charontoidea are similar to the older groups Pulvillata and Apulvillata, respectively, as conceived by Quintero (1986). However, these names were abandoned by Weygoldt (1996, 2000) and Harvey (2003), so we follow Weygoldt (1996) and Harvey (2003) in naming Phrynichidae + Phrynidae as Phrynoidea, and Charontoidea for Charinidae + Charontidae.

**Figure 3.**
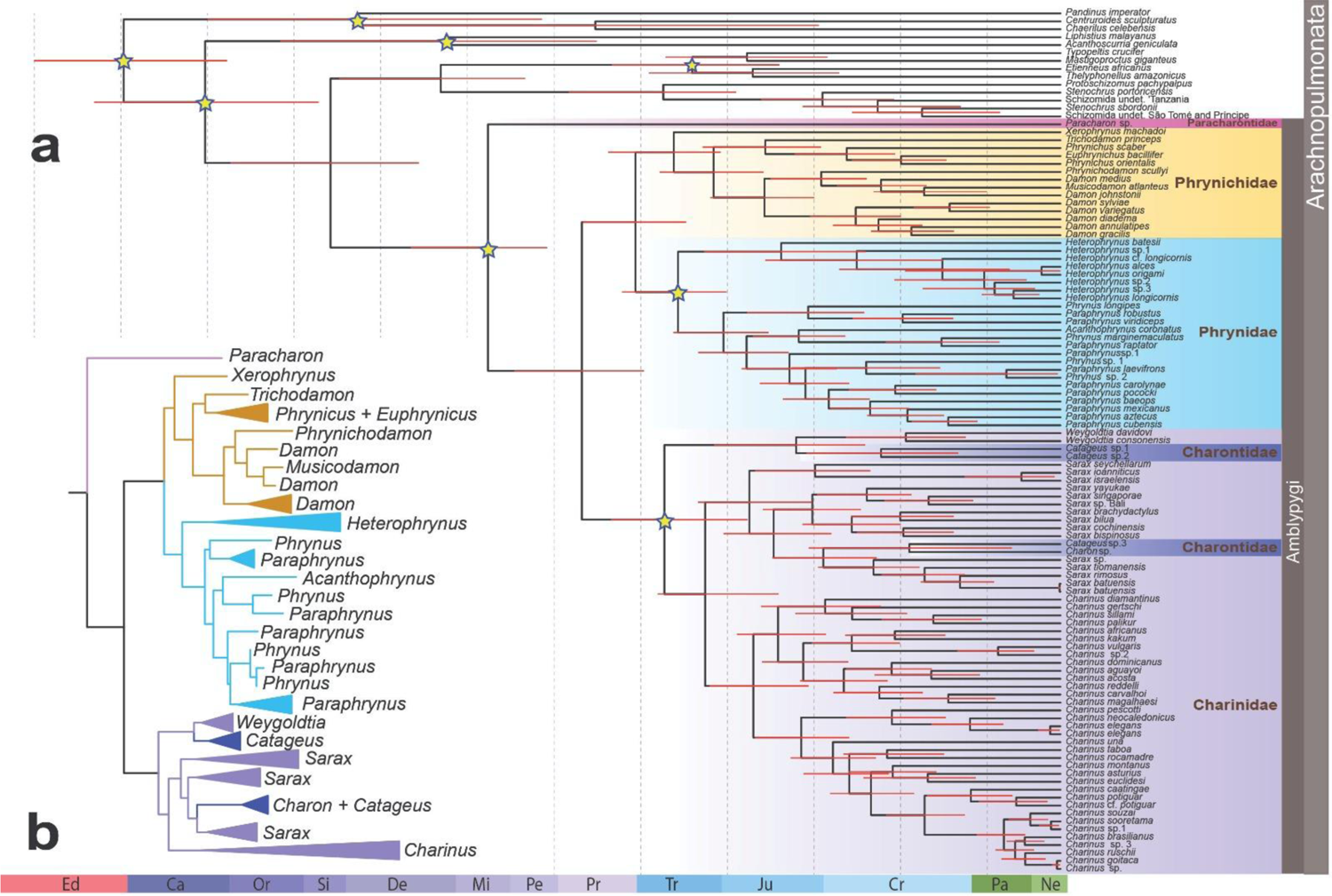
a) Divergence time estimation for Amblypygi, based on five outgroup and three ingroup node calibrations marked with a star. Data matrix is derived from a fusion of UCEs (40% occupancy) and six Sanger loci. b) Simplified maximum likelihood tree of the UCE and Sanger dataset, showing intergeneric relationships.

Topologies of phrynichid genera in analyses of UCE matrices (Fig. 2; Supplementary Fig. S1) were congruent with the morphological cladistic analysis of Miranda et al. (2018). In Phrynidae, monophyly of each of the two subfamilies Heterophryninae and Phryninae was recovered. *Charon*, the type genus of Charontidae, was recovered as a sister group to (10% and 40% occupancy), or nested within (25% occupancy matrix), Charinidae (Fig. 2; Supplementary Fig. S1), contradicting Weygoldt (1996) and Garwood et al. (2017), who recovered Charinidae as sister-group to the remaining Euamblypygi taxa.

Analyses of the 109-terminal UCE + Sanger loci dataset also recovered *Paracharon* sp. as the sister group to Euamblypygi (Fig. 3), a clade formed by Phrynoidea and Charontoidea. In Phrynichidae, *Xerophrynus* was recovered sister to all other Phrynichidae genera, followed by the divergence of the reciprocally monophyletic subfamilies Phrynichinae and Damoninae. The topology for Phrynichinae accords with morphological analyses by Weygoldt (1996) and Miranda et al. (2018), with *Trichodamon* sister to *Phrynichus* + *Euphrynichus*. In Damoninae, *Damon* was found polyphyletic, with *D. medius* and *D. johnstoni* nested in a clade with *Musicodamon* and *Phrynichodamon*. Phrynidae was recovered in two main branches, Heterophryninae and Phryninae. *Phrynus* and *Paraphrynus* were recovered as polyphyletic.

Within Charontoidea, Charontidae (represented by a single *Charon* and three *Catageus*; Miranda et al. 2018) was recovered as polyphyletic, and all four exemplars were nested within Charinidae with robust support. Two species identified as *Catageus* by Weygoldt (in Arabi et al. 2011) were recovered as the sister group to *Weygoldtia*, and together formed the sister group to the remaining Charontoidea. The third species identified as *Catageus* by Weygoldt (in Arabi et al. 2011) was recovered in a clade with *Charon* that was nested within *Sarax*. The non-monophyly of various genera support the need for systematic revision of shallow-level taxa, as exemplified by recent changes implemented within the phylogeny of Charinidae (Miranda et al. 2022).

Our analyses thus revealed phylogenetic relationships that conflicted with morphological interpretations with respect to the position of Charinidae (unambiguously forming a clade with Charontidae, *contra* Weygoldt 2000; Garwood et al. 2017), for all matrices analyzed, regardless of occupancy and taxon composition. When support for (Charinidae +_Charontidae) was assessed, the likelihood mapping results strongly favor the placement of Charinidae and Charontidae as sister groups (78.5% of quartets; Fig. 2b). From the unfiltered UCE data set, 2.1% of the sampled quartets were uninformative.

### Molecular dating

Divergence time estimation using five outgroup and three ingroup fossil calibrations in MCMCTree recovered a 330 Mya age for the crown group of Amblypygi (95% highest posterior density [HPD] interval: 374-295 Mya), with diversification of family-level taxa occurring in the late Permian and Triassic (Fig. 3). Removal of *Paracharon* resulted in node age overestimates for the deepest nodes in Ambypygi, but negligible effects on shallow nodes (Supplementary Fig. S2). The estimated age of Charinidae accorded closely with previous estimates based on two fossil calibrations and a reduced representation of Amblypygi higher-level diversity (Miranda et al. 2022).

By comparison to MCMCTree, divergence time estimation using a penalized likelihood approach with LSD2 exhibited a more pronounced impact upon the removal of *Paracharon*. When *Paracharon* sp. was removed and a node calibration from the oldest known whip spider fossil was instead applied to the most recent common ancestor of Euamblypygi, the resulting analysis recovered a 314 Mya age for crown group Euamblypygi (95% HPD interval: 353-314 Mya), and divergences of the family-level taxa in the Jurassic. All node ages were overestimated by LSD2 when *Paracharon* sp. was excluded, with the magnitude of overestimation correlating with phylogenetic depth (Supplementary Fig. S2). This result suggests that the sampling of Paracharontidae strongly influences divergence time estimation within Amblypygi, as a function of estimation method.

To compare the relative influence of denser sampling of extant lineages versus the use of ingroup fossil calibrations, we performed a third analysis wherein we retained all terminals, but removed the three ingroup fossil calibrations. Divergence time analyses using only outgroup node calibrations resulted in broader variance (HPD intervals) for all node age estimates of interest, regardless of the use of MCMCTree or LSD2 (Fig. 4), but median age estimates did not significantly change as a result of removing the ingroup fossil calibrations.

**Figure 4.**
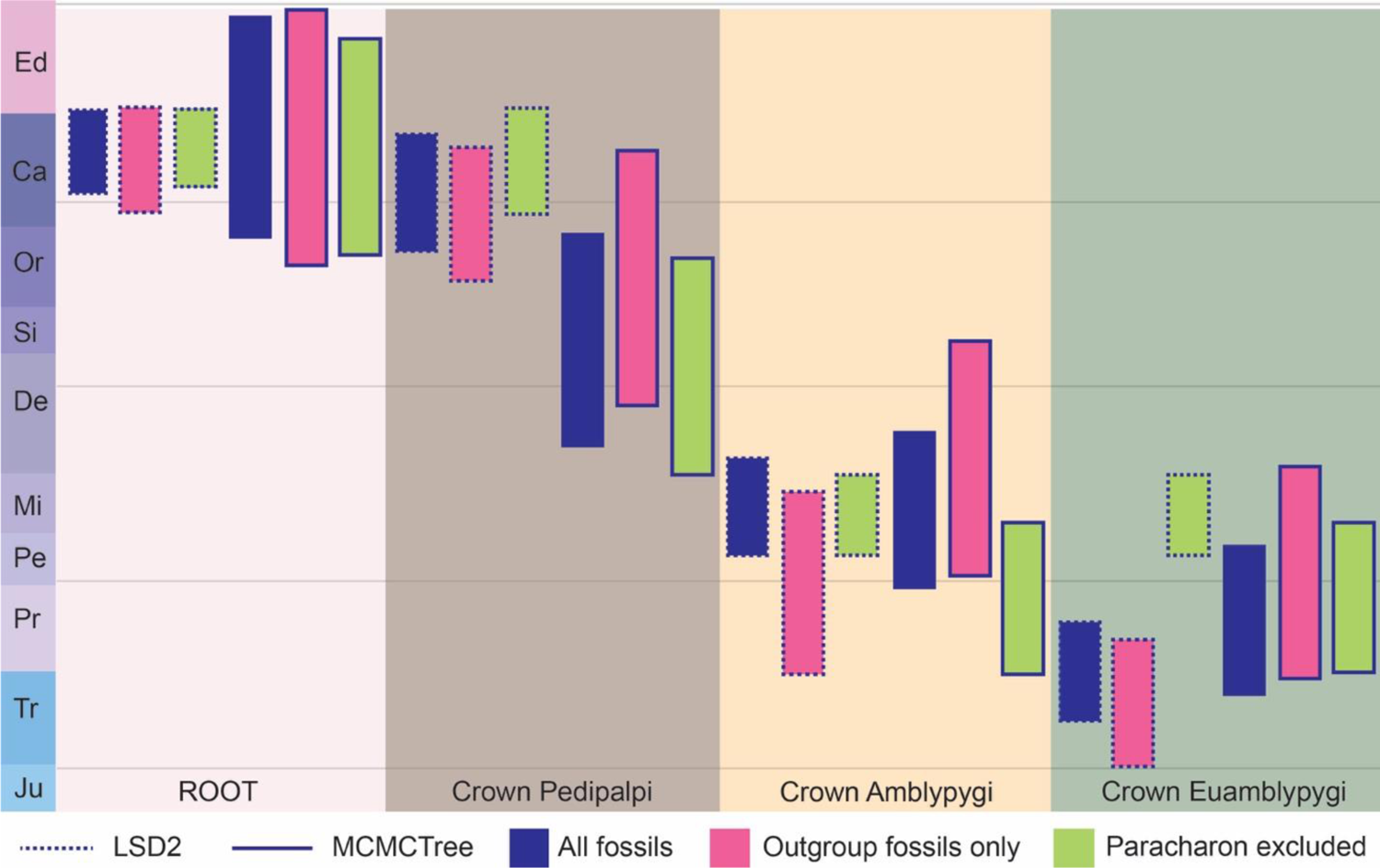
Divergence time estimates optimized by MCMCTree and LSD2 using fossil-based node calibrations. Conventions of HPD intervals follow the bottom legend.

## Discussion

Considerable strides have been achieved in deciphering the relationships of chelicerate arthropods in the past decade. An emphasis on denser taxonomic sampling, together with the availability of new sequencing technologies, has revolutionized modern investigations of relationships within poorly studied groups, such as mites, pseudoscorpions, and sea spiders (e.g., Klimov et al. 2018; Benavides et al. 2019; Ballesteros et al. 2021). Such datasets have facilitated greater precision in hypothesis-testing with regard to diversification dynamics, biogeographic history, and both morphological and molecular evolution (e.g., Giribet et al. 2012; Bond et al. 2014; Wheeler et al. 2017; Benavides et al. 2019; Santibáñez-López et al. 2022). Surprisingly, there remain a handful of higher level taxa within arthropods for which molecular phylogenies still do not exist, as exemplified by Amblypygi. In this specific case, the sister group to the remaining whip spiders long consisted of a single elusive relict species that had not been rediscovered for over a century, effectively barring any sequencing-based efforts to infer higher-level phylogeny. The phylogeny of Amblypygi was last assessed in an analytical framework using morphological characters by Weygoldt (1996). Thereafter, phylogenetic works focused on small species-groups (Prendini et al., 2005; Seiter et al., 2020), fossil placements (Garwood at al., 2017), or the systematics of individual families (Miranda et al., 2018, 2021); relationships across the order were never reinvestigated. Here, our rediscovery of a long-lost relict breaks an erstwhile phylogenetic impasse, greatly expands the known range of the relictual lineage Paracharontidae, and significantly improves the precision of molecular dating efforts.

Our analyses of whip spider relationships recovered a well-resolved topology that placed Paracharontidae as the sister group of the remaining Amblypygi, consistent with the interpretations of Weygoldt (1996) and Garwood et al. (2017). Paracharontids closely resemble Carboniferous fossils of Amblypygi, such as *Weygoldtina anglica* (315 Ma). The articulation of the paracharontid pedipalp is parallel to the body axis, in contrast to the remaining Amblypygi. This feature is shared by the Paleozoic fossil whip spiders, as well as by non-whip spider arachnopulmonates, such as Schizomida and spiders. All analyses additionally recovered the clades Phrynidae + Phrynichidae, as well as Charinidae + Charontidae with robust support. These results are partly congruent with each of the preceding morphological analyses of Amblypygi (Quintero 1986; Weygoldt 1996; Garwood et al. 2017).

The undescribed species of *Paracharon* that we identified from the Neotropics is morphologically similar to *Paracharon caecus* from Guinea Bissau, including in its extreme reduction of eyes and its troglobitic habitus, which likely reflects a case of convergence (Hansen 1921). Together with the Indian amber species *Paracharonopsis cambayensis*, as well as the Nearctic Carboniferous taxa (*Weygoldtina*; *Graeophonus*), these localities suggest a broad former distribution of Paleoamblypygi that underwent significant range contraction. Given the comparatively large size and visibility of whip spiders, we postulate that the remaining species of Paracharontidae may be restricted to caves and other microrefugia of Tropical Gondwana.

Beyond testing the validity of phylogenetic relationships inferred on the basis of morphology, there are two critical benefits to phylogenetic analyses that hinge upon sampling phylogenetic relicts with molecular data. First, we demonstrate that the omission of relicts has a significant impact on node dating approaches. When *Paracharon* sp. is artificially excluded from the dataset, the next available crown group node is treated as the most recent common ancestor of Amblypygi. The effect of this incorrect calibration is that the crown age of Amblypygi is overestimated, particularly by penalized likelihood approaches. This overestimation of node ages has downstream implications for any hypothesis-testing that relies upon divergence time estimation (namely, historical biogeography; Miranda et al. 2022). Second, phylogenetic relicts such as *Paracharon* have the effect of breaking branches that subtend crown group taxa (Inward et al. 2007; Soltis et al. 2008). The sampling of such basally branching groups is critical to phylogenetic accuracy when unstable relationships are driven by heterogeneous evolutionary rates across taxa. In the case of chelicerate phylogeny, a recent work showed that the strategy of densely sampling the basal nodes of an unstable taxon outperformed filtering by taxon occupancy, filtering by evolutionary rate, algorithmic approach to tree reconstruction, and use of site heterogeneous models, as predictors of phylogenetic accuracy (Ontano et al. 2021).

Our results, therefore, underscore the imperative for biodiversity discovery and continued campaigns to survey and document the diversity of tropical invertebrates, with the goal of finding taxa that represent phylogenetically and evolutionarily significant groups, including relicts (Griswold et al. 2012; Cruz-López et al. 2016; Aharon et al. 2019). Future efforts to advance invertebrate systematics must emphasize the completion of higher-level phylogenies for dark parts of the Tree of Life, in tandem with recognition of the scientific value of depauperate groups of phylogenetic significance.

## Acknowledgements

This work is dedicated to the legacy of Peter Weygoldt. Access to natural history collections was provided by Jairo Moreno González, Lorenzo Prendini, Gonzalo Giribet, Lauren Esposito, Joachim Holstein, and Nikolaj Scharff, and access to recently-collected material from Australia was provided by Kieran Aland. Siegfried Huber provided the photograph of *Damon medius*. Sample preparation for sequencing was performed at the Laboratory of Analytical Biology, Smithsonian Institution, Washington, DC; and at the BioTechnology Center, University of Wisconsin-Madison. Scientific computation was performed on the Smithsonian High Performance Cluster (SI/HPC), Smithsonian Institution, High Performance Computing Cluster at The George Washington University, Research Technology Services, and at the Biotechnology Resource Center, University of Wisconsin-Madison.

