## Supplementary Table S1 for "The rediscovery of a relict unlocks the first global phylogeny of whip spiders (Amblypygi)"

**Supplementary Table S1.** Extant diversity of phylogenetic relicts within Chelicerata that constitute sister groups to extant orders.

| Relict | Parent | Relict species | Non-relict sister species | Parent species | Proportion |
|---|---|---:|---:|---:|---:|
| Mesothelae | Araneae | 182 | 49818 | 50000 | 0.00364 |
| Cyphophthalmi | Opiliones | 180 | 6339 | 6519 | 0.0276116 |
| Austrodecidae | Pycnogonida | 75 | 1255 | 1330 | 0.05639098 |
| Opilioacariformes | Parasitiformes | 37 | 12347 | 12384 | 0.00298773 |
| Protoschizomidae | Schizomida | 11 | 253 | 264 | 0.04166667 |
| Prokoeneniidae | Palpigradi | 7 | 75 | 82 | 0.08536585 |
| Nanorchestidae | Acariformes | 2 | 42231 | 42233 | 4.7356E-05 |
| Paracharontidae | Amblypygi | 1 | 259 | 260 | 0.00384615 |
