## Supplementary Material for "The rediscovery of a relict unlocks the first global phylogeny of whip spiders (Amblypygi)"

**This PDF file includes:**

Supplementary Methods
Supplementary Figures S1, S2
Supplementary Tables S1-S4

### Supplementary Methods

*Library preparation for ultraconserved element sequencing*

The homogenate from DNA extraction was incubated at 55 °C overnight and purified following the manufacturer's protocol. The DNA extractions were quantified using high sensitivity Qubit fluorometry (Life Technologies, Inc.) and quality checked using gel electrophoresis on a 1.5% agarose gel.

Depending on prior degradation and quality of the DNA, between 7 and 100 ng of DNA were sheared between 10 and 60 s (amp=25%, pulse=10–10 seconds, to a target size of approximately 250–600 bp) by sonication (Q800R, Qsonica Inc.). Sheared DNA was dried completely and rehydrated to the required input volume (13 µL) and used as input for DNA library preparation (Kapa Hyper Prep Library kit, Kapa Biosystems). After ligation of universal stubs (Faircloth and Glenn, 2012), a 0.8× SPRI bead clean was done (Kapa Pure Beads, Kapa Biosystems) on a Wafergen Apollo liquid handler (Wafergen Biosystems), resulting in 30 µL of post-ligation library. For adapter ligation, we used TruSeq-style adapters (Faircloth and Glenn, 2012). PCR conditions were as follows: 15 µL post ligation library, 25 µL HiFi HotStart polymerase (Kapa Biosystems), 2.5 µL each of Illumina TruSeq-style i5 and i7 primers, and 5 µL double-distilled water (dd$H_2O$). We used the following thermal protocol (Kapa Biosystems): 98 °C for 45 s; 13 cycles of 98 °C for 15 s, 65 °C for 30 s, 72 °C for 60 s, and final extension at 72 °C for 5 m. PCR cleanup was done with a 0.8 X SPRI bead clean (Kapa Pure Beads, Kapa Biosystems) on a Wafergen Apollo (TaKaRa Bio Inc. USA) with a final library volume of 20 µL. Following clean-up, libraries were divided into enrichment pools containing eight libraries combined at equimolar ratios with final concentrations of 137–184 ng/µL.

Hybridization reactions were incubated for 24 h at 65 °C, subsequently, all pools were bound to streptavidin beads (MyOne C1; Life Technologies), and washed. We combined 15 µL of streptavidin bead-bound, washed, enriched library with 25 µL HiFi HotStart Taq (Kapa Biosystems), 5 µL of Illumina TruSeq primer mix (5 µM forward and reverse primers) and 5 µL of ddH2O. Post-enrichment PCR used the following thermal profile: 98 °C for 45 s; 18 cycles of 98 °C for 15 s, 60 °C for 30 s, 72 °C for 60 s; and a final extension of 72 °C for 5 m. We purified the resulting reactions using 1X bead clean using Kapa Pure Beads (Kapa Biosystems), and resuspended the enriched pools to total 22 µL.

We then quantified pools using qPCR library quantification (Kapa Biosystems) with two serial dilutions of each pool (1:100,000, 1:1,000,000), assuming an average library fragment length of 600 bp. Based on the size-adjusted concentrations estimated by qPCR, we combined all pools at an equimolar concentration of 30 nM, and size selected for 250–600 bp with a BluePippin (SageScience). We sequenced the pooled libraries in a single lane of a paired-end run on an Illumina HiSeq 2500 (2x150bp rapid run) at the University of Utah Huntsman Cancer Institute, or an Illumina NovaSeq 6000 (2x150 bp) at the University of Wisconsin-Madison.

**Supplementary Figure 1.** Phylogenetic relationships of Amblypygi reconstructed using UCE data at occupancies; a) 10%; b) 25%; c) 40%.

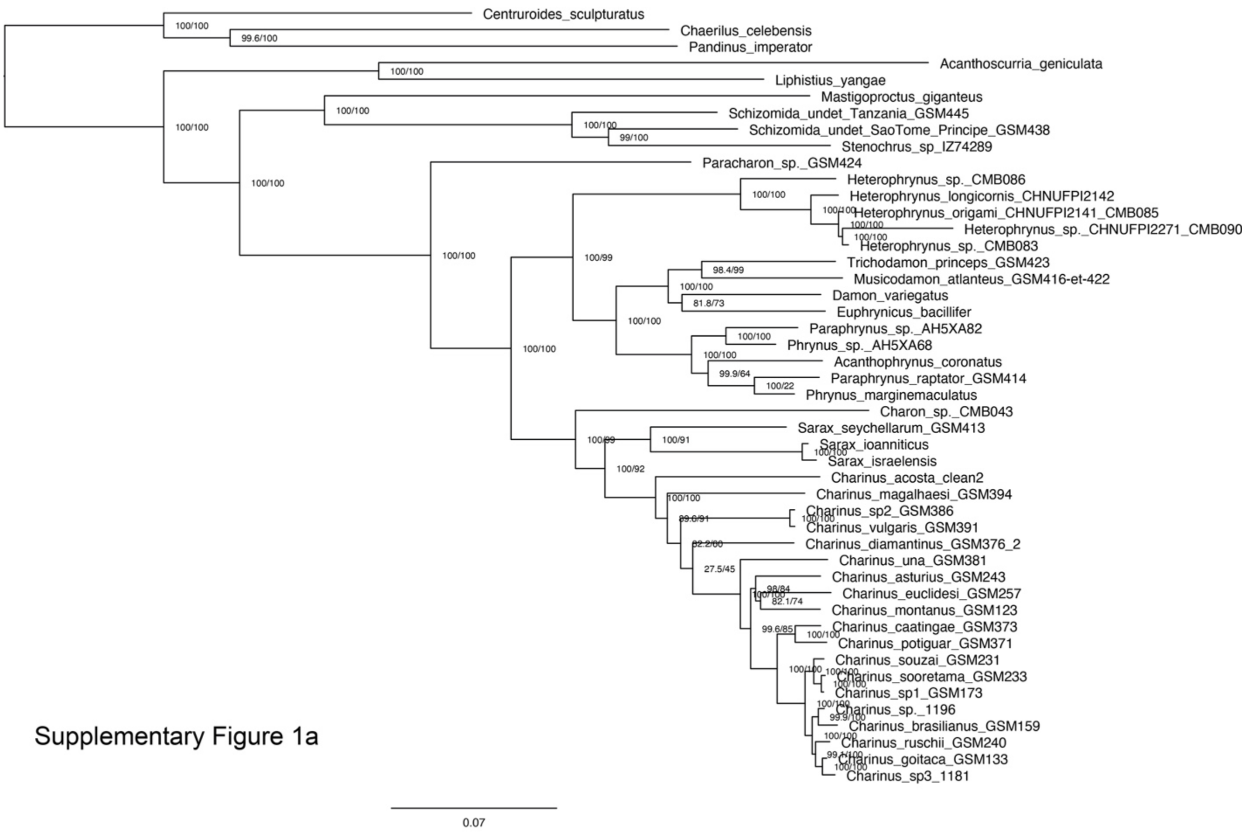

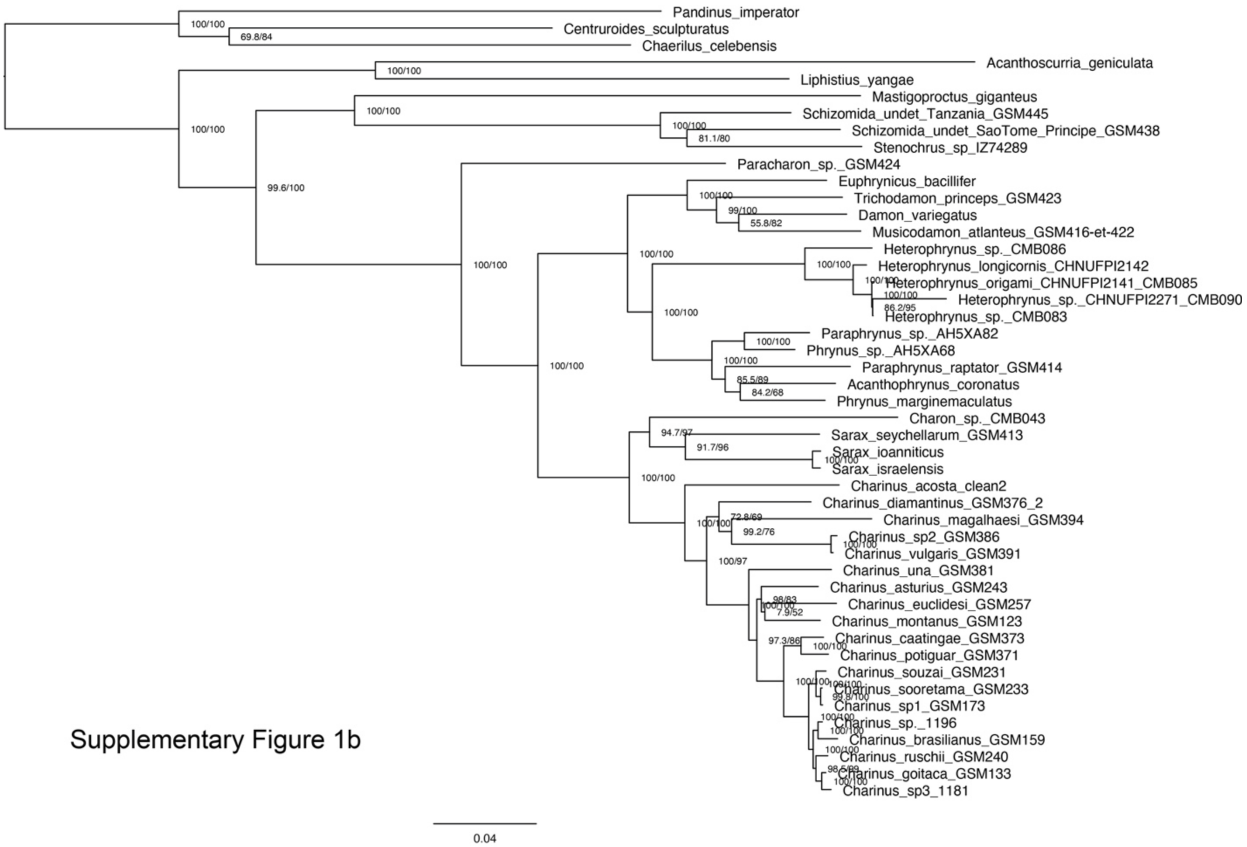


Supplementary Figure 1b

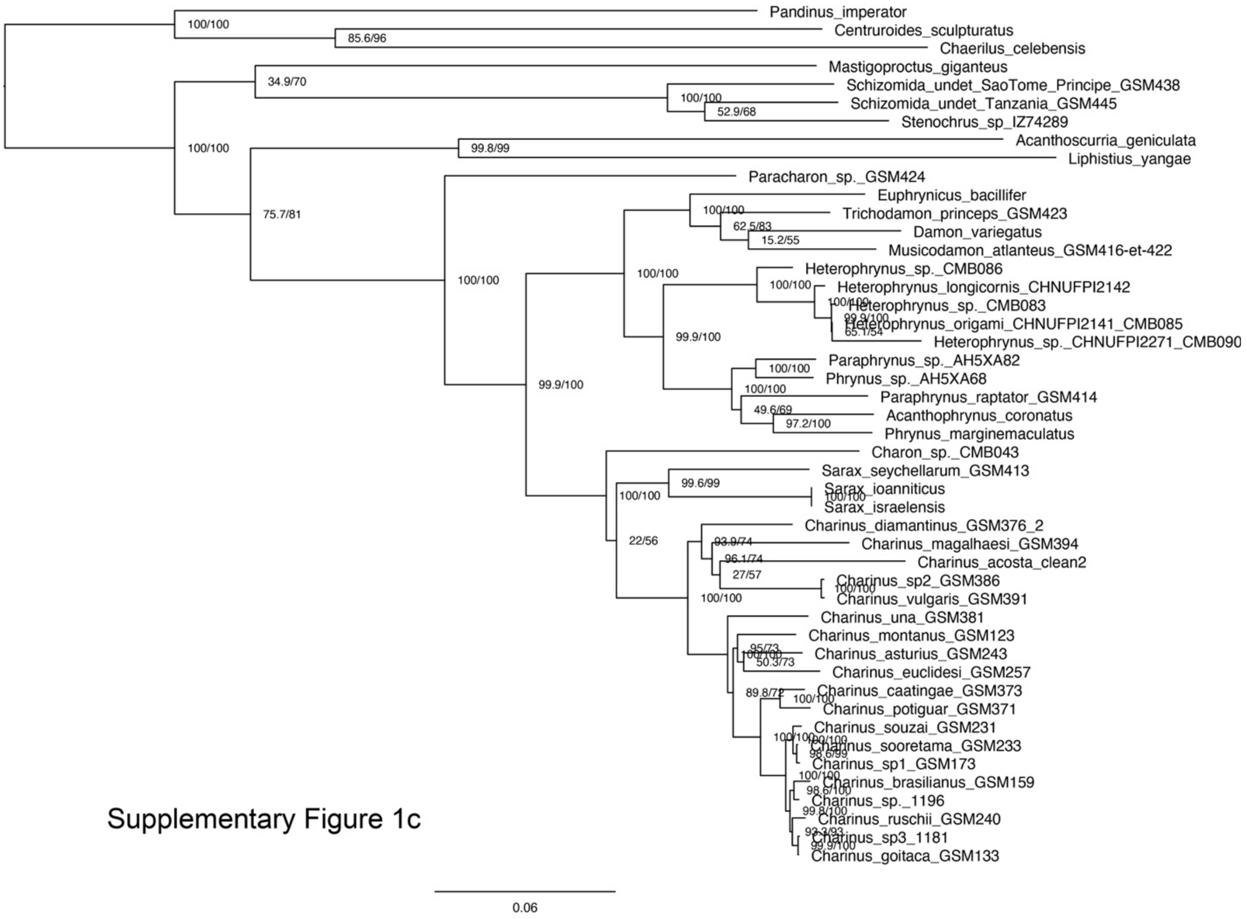


Supplementary Figure 1c

**Supplementary Figure 2.** Median node ages inferred by MCMCTree (top) and LSD2 (bottom), upon retention (purple) or exclusion (green) of Paracharontidae.

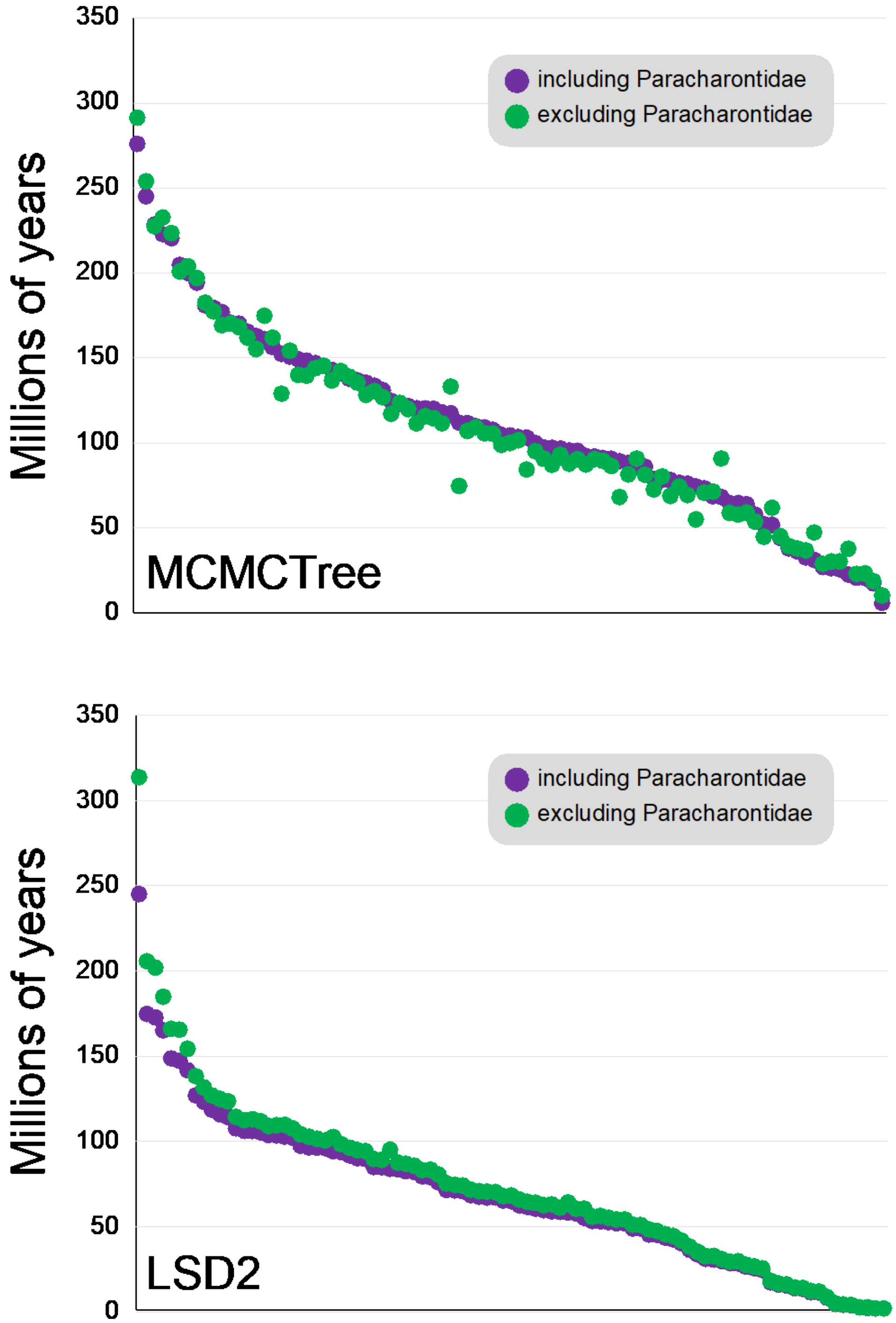

**Supplementary Table S1.** Extant diversity of phylogenetic relicts within Chelicerata that constitute sister groups to extant orders.

| Relict | Parent | Relict species | Non-relict sister species | Parent species | Proportion |
|---|---|---|---|---|---|
| Mesothelae | Araneae | 182 | 49818 | 50000 | 0.00364 |
| Cyphophthalmi | Opiliones | 180 | 6339 | 6519 | 0.0276116 |
| Austrodecidae | Pycnogonida | 75 | 1255 | 1330 | 0.05639098 |
| Opilioacariformes | Parasitiformes | 37 | 12347 | 12384 | 0.00298773 |
| Protoschizomidae | Schizomida | 11 | 253 | 264 | 0.04166667 |
| Prokoeneniidae | Palpigradi | 7 | 75 | 82 | 0.08536585 |
| Nanorchestidae | Acariformes | 2 | 42231 | 42233 | 4.7356E-05 |
| Paracharontidae | Amblypygi | 1 | 259 | 260 | 0.00384615 |

**Supplementary Table S2.** Locality and accession data for terminals included in the study. Nuclear histone 3, H3; mitochondrial cytochrome c oxydase subunit I, COI; nuclear 28S rDNA, 28S; nuclear 18S rDNA, 18S; mitochondrial 16S rDNA, 16S; mitochondrial 12S rDNA, 12S

| Voucher codes | **Sequencing sucess - GenBank Acc.** | **Chimera** | **Taxon, author** | **Locality** |
|---|---|---|---|---|
| | SRX815750 | No | *Pandinus imperator* (Koch, 1842) | Africa |
| | SRR3932806 | No | *Centruroides sculpturatus* (Wood, 1863) | California, Arizona, New Mexico |
| | SRX815898 | No | *Chaerilus celebensis* (Pocock, 1894) | Indonesia, Luwu |
| | SRX450965 | No | *Liphistius malayanus* Abraham, 1923 | Malaysia |
| | GCA_000661875 | No | *Acanthoscurria geniculata* (Koch, 1841) | Brazil |
| ARALP000154/ ARALP000069 | H3-KY018462 COI-KY017961 28S-KY017406 18S-KY016748 16S-KY016154 12S-KY015604 | No | *Typopeltis crucifer* Pocock, 1894 | Japan, Taiwan |
| MNHN-JAA17/ ARALP000013/ LP 3509 | H3-FJ862878 28S-JN018408 18S-AF005446 16S-KY016153 12S-KY509342 | **Yes** | *Mastigoproctus giganteus* (Pocock, 1894) | Mexico, USA |
| MNHN-JAC98 | COI-JN018215 (part) | **Yes** | *Mastigoproctus giganteus (Pocock, 1894)* | Mexico, USA |
| ARALP000174 | COI-KY017960 (part) | **Yes** | *Mastigoproctus giganteus (Pocock, 1894)* | Mexico, USA |
| ARALP000172 | COI-KY017959 18S-KY016746 16S-KY016152 | No | *Etienneus africanus* (Hentschel, 1899) | Gambia, Senegal |
| MCZ IZ-129084/ TH01 | COI-KY573153 28S-KP276369 18S-KY573511 | No | *Thelyphonellus amazonicus* Butler, 1872 | Brazil, Surinam |
| AMNH-LP14518 | COI-MN103943 28S-MN108254 12S-MN108252 | No | *Protoschizomus pachypalpus* (Rowland, 1973) | Mexico |
| AMNH-LP3755/ AMNH-LP6014/ FL47.2 | COI-MN097762 18S-AF005444 16S-MN999751 12S-KY573465 | No | *Stenochrus portoricensis* (Chamberlin, 1922) | Central America, South America, North America, |
| CASENT 9060741 | UCE, this study | No | Schizo Tanzania | Tanzani, Eastern Arc Mountains |

| | | | | |
|---|---|---|---|---|
| ARALP000138/ ARALP000003/ AMNH-LP3757 | H3-KY018263 COI-KY017755 28S-KY017130 18S-KY509361 12S-MK849658 | No | *Stenochrus sbordonii* (Brignoli, 1973) | Mexico |
| CASENT 9060721 | UCE, this study | No | SchizoSaoTome_Principe | Sao Tome and Principe |
| MNRJ | UCE, this study | No | *Paracharon*sp. | Colombia |
| AMCC 124710 | H3-AY829961 28S-AY829921 18S-AY829900 16S-AY829879 12S-AY829860 | No | *Xerophrynus machadoi* (Fage, 1951) | Angola, Namibia |
| ARP-2015 | 28S-KP276346 18S-KP276426 | No | *Trichodamon princeps* (Mello-Leitão, 1935) | Brazil |
| AMCC 124712 | H3-AY829962 COI-AY829942 28S-AY829922 18S-AY829901 16S-AY829880 12S-AY829861 | No | *Phrynichus scaber* (Gervais, 1844) | Mauritius, Seychelles |
| AMCC 124711 | H3-AY829963 COI-AY829943 28S-AY829923 18S-AY829902 16S-AY829881 12S-AY829862 | No | *Euphrynichus bacillifer* (Gerstaecker, 1873) | East Africa |
| MNHN-JAA1/ MNHN-JAB13 | COI-JN018195 28S-JN018330 18S-JN018233 | No | *Phrynichus orientalis* Weygoldt, 1998 | Cambodia, Malaysia, Thailand, Vietnam |
| AMCC 124713 | H3-AY829965 COI-AY829944 28S-AY829925 18S-AY829904 16S-AY829883 12S-AY829864 | No | *Phrynichodamon scullyi* (Purcell, 1902) | South Africa |
| MNHN-JAB60/ AMCC 124715 | 28S-JN018405 18S-JN018308 16S-AY829884 | **Yes** | *Damon medius* (Herbst, 1797) | Africa |
| MNHN-JAB60 | COI-JN018192 (part) | **Yes** | *Damon medius* (Herbst, 1797) | Africa |
| AMCC 124715 | COI-AY829945 (part) | **Yes** | *Damon medius* (Herbst, 1797) | Africa |
| AMCC 124714 | H3-AY829964 28S-AY829924 18S-AY829903 16S-AY829882 12S-AY829863 | No | *Musicodamon atlanteus* Fage, 1939 | Algeria, Morocco |
| MNHN-JAD13 | COI-JN018115 28S-JN018329 18S-JN018232 | No | *Damon johnstoni* (Lydekker, 1879) | Africa |
| AMCC 124724 | COI-AY829952 28S-AY829933 18S-AY829912 16S-AY829891 12S-AY829872 | No | *Damon sylviae* Prendini, Weygoldt & Wheeler, 2005 | Namibia |
| AMCC 124730/ AMCC 124725/ AMCC124727 | H3-AY829979 COI-AY829958 28S-AY829939 18S-AY829918 16S-AY829892 12S-AY829874 | No | *Damon variegatus* (Perty, 1834) | Africa |
| ARALP000030/ AMCC 124716/ AMCC 124717 | H3-KY018357 COI-AY829946 28S-KY017271 18S-AY829907 16S-AY829886 12S-AY829866 | No | *Damon diadema* (Simon, 1876) | Africa |
| AMCC 124718/ AMCC 124720/ AMCC 124719 | COI-AY829948 28S-AY829931 18S-AY829910 16S-AY829888 12S-AY829869 | No | *Damon annulatipes* (Wood, 1869) | South Africa |

| | | | | |
|---|---|---|---|---|
| AMCC 124722/ MNHN-JAB38/ AMCC 124721 | H3-AY829981 28S-JN018328 18S-AY829920 16S-AY829899 12S-AY829878 | **Yes** | *Damon gracilis* Weygoldt, 1998 | Angola, Namibia |
| MNHN-JAB38 | COI-JN018114 (part) | **Yes** | *Damon gracilis* Weygoldt, 1998 | Angola, Namibia |
| AMCC 124721 | COI-AY829959 (part) | **Yes** | *Damon gracilis* Weygoldt, 1998 | Angola, Namibia |
| Hb-B-CRdM-UFMG15422 | COI-MT899143 (part) | No | *Heterophrynus batesii* (Butler, 1873) | Brazil |
| CMB 086 | UCE, this study | No | *Heterophrynus* sp.1 | Brazil, Amazonas, Guajará |
| AMCC_LP_3830/ MNHN-JAD3 | COI-MW410162 28S-JN018333 18S-JN018236 | No | *Heterophrynus* cf. *longicornis* (Butler, 1873) | French Guiana: Approuague-Kaw,Kaw mountains |
| MNHN-JAA81 | COI-JN018118 28S-JN018332 18S-JN018235 | No | *Heterophrynus alces* (Pocock, 1902) | Brazil, Guyana, Surinam |
| CHNUFPI 2141 | UCE, this study | No | *Heterophrynus origami* Chirivi-Joya, Moreno-González & Fagua, 2020 | Brazil, Rondonia, Jamari |
| CMB 083 | UCE, this study | No | *Heterophrynus* sp.2 | Brazil, Amazonas, Canutama |
| CMB 090 | UCE this study | No | *Heterophrynus* sp.3 | Brazil, Amazonas, Canutama |
| ARALP000188/ LP 1483/ ARALP000107 | H3-KY018358 COI-MT040911 28S-KY017272 18S-KY509344 16S-MF806089 12S-MF806134 | No | *Phrynus longipes* (Pocock, 1894) | Dominican Republic, Haiti, Puerto Rico, Virgin Islands |
| 13872 | COI-MT738756 28S-MT734793 18S-MT734777 16S-MT734766 12S-MT753022 | No | *Paraphrynus robustus* (Franganillo, 1931) | Cuba |
| 13881 | COI-MT738757 28S-MT734794 18S-MT734778 16S-MT734767 | No | *Paraphrynus viridiceps* (Pocock, 1894) | The Bahamas, Cuba |
| AMCC_LP_16891/ AMCC_LP_5374/ AMCC_LP_16893 | COI-MW410159 28S-MW411456 16S-MW411542 | **Yes** | *Acanthophrynus coronatus* (Butler, 1873) | Mexico, USA |
| AMCC_LP_8672 | 18S-MW411509 (part) | **Yes** | *Acanthophrynus coronatus* (Butler, 1873) | Mexico, USA |
| AMCC_LP_16886 | 18S-MW411499 (part) | **Yes** | *Acanthophrynus coronatus* (Butler, 1873) | Mexico, USA |
| AMCC_LP_8672 | 18S-MW411485 (part) | **Yes** | *Acanthophrynus coronatus* (Butler, 1873) | Mexico, USA |
| AMCC_LP_5374 | 12S-MW411482 (part) | **Yes** | *Acanthophrynus coronatus* (Butler, 1873) | Mexico, USA |
| AMCC_LP_16886 | 12S-MW411499 (part) | **Yes** | *Acanthophrynus coronatus* (Butler, 1873) | Mexico, USA |

| | | | | |
|---|---|---|---|---|
| 14072 | 18S-MT734779 16S-MT734768 12S-MT753024 | **Yes** | *Phrynus marginemaculatus* (C. L. Koch, 1841) | The Bahamas, Bermuda, Cayman Islands, Cuba, Dominican Republic, Haiti, Jamaica, Puerto Rico, USA |
| 80122 | COI-MH478994 (part) | **Yes** | *Phrynus marginemaculatus* (C. L. Koch, 1841) | The Bahamas, Bermuda, Cayman Islands, Cuba, Dominican Republic, Haiti, Jamaica, Puerto Rico, USA |
| 14072 | COI-MT738758 (part) | **Yes** | *Phrynus marginemaculatus* (C. L. Koch, 1841) | The Bahamas, Bermuda, Cayman Islands, Cuba, Dominican Republic, Haiti, Jamaica, Puerto Rico, USA |
| AMNH | UCE, this study | No | *Paraphrynus raptator* (Pocock, 1902) | Honduras, Dept. Comayagua, Meambar |
| USNM AH5XA82 | UCE, this study | No | *Paraphrynus* sp.1 | Costa Rica, Heredia, Puerto Viejo de Sarapiqui |
| USNM AH5XA88 | UCE, this study | No | *Phrynus* sp.1 | Costa Rica, Heredia |
| AMCC_LP_6145 | COI-MW410163 28S-MW411480 18S-MW411515 16S-MW411544 12S-MW411573 | No | *Paraphrynus laevifrons* (Pocock, 1894) | Costa Rica, Puntarenas Province, Fundacion Neotropica |
| USNM AH5AX88 | UCE, this study | No | *Phrynus* sp.2 | Costa Rica, Heredia |
| 14444 | COI-MT738750 28S-MT734787 18S-MT734771 16S-MT734760 12S-MT753016 | No | *Paraphrynus carolynae* de Armas, 2012 | Mexico, USA |
| 2091 | COI-MT738753 28S-MT734790 18S-MT734774 16S-MT734763 12S-MT753019 | No | *Paraphrynus pococki* Mullinex, 1975 | Mexico |
| 8667 | COI-MT738749 28S-MT734786 18S-MT734770 12S-MT753015 | No | *Paraphrynus baeops* Mullinex, 1975 | Mexico |
| 15431 | COI-MT738752 28S-MT734789 18S-MT734773 16S-MT734762 12S-MT753018 | No | *Paraphrynus mexicanus* (Bilimek, 1867) | Mexico |
| 2096 | COI-MT738748 28S-MT734785 18S-MT734769 16S-MT734759 12S-MT753014 | No | *Paraphrynus aztecus* (Pocock, 1894) | Mexico |
| 13883 | COI-MT738751 28S-MT734788 18S-MT734772 16S-MT734761 12S-MT753017 | No | *Paraphrynus cubensis* (Quintero, 1983) | Cuba |
| LP 11377/ LP 11375 | COI-MT040904 28S-KY509389 18S-KY509339 16S-MF806085 12S-MF806128 | No | *Weygoldtia davidovi* (Fage, 1946) | Cambodia, Laos, Vietnam |
| LP 11269 | COI-MT040912 28S-KY509395 18S-KY509345 16S-MF806090 12S-MF806135 | No | *Weygoldtia consonensis* Miranda, Giupponi, Prendini & Scharff, 2021 | Con Son Island |
| MNHN-JAA93 | COI-JN018111 28S-JN018325 18S-JN018228 | No | *Catageus* sp.1 | Unknown |
| MNHN-JAB1 | COI-JN018112 28S-JN018326 18S-JN018229 | No | *Catageus* sp.2 | Unknown |
| LP 9074 | COI-MT040928 28S-KY509413 18S-KY509363 16S-MF806105 MF806153 | No | *Sarax seychellarum* (Kraepelin, 1898) | Seychelles |
| LP 13394/ LP2843 | COI-MT040910 28S-KY509393 18S-KY509343 16S-MF806088 12S-MF806133 | No | *Sarax ioanniticus* (Kritscher, 1959) | Egypt, Greece, Jordan, Israel, Italy, Turkey |

| | | | | |
|---|---|---|---|---|
| | SRX8887651-SRX8887663 | No | *Sarax israelensis* (Miranda, Aharon, Gavish-Regev, Giupponi & Wizen, 2016) | Israel |
| LP 12169/ LP 12123 | COI-MT040945 28S-KY509430 18S-KY509380 16S-MF806116 12S-MF806167 | No | *Sarax yayukae* Rahmadi, Harvey & Kojima, 2010 | Sabah, Sarawak - Malaysia |
| LP 4761 | COI-MT040931 28S-KY509416 18S-KY509366 16S-MF806107 12S-MF806156 | No | *Sarax singaporae* (Gravely, 1911) | Singaporae |
| LP 11594 | COI-MT040925 28S-MF806103 18S-MF806150 16S-KY509410 12S-KY509360 | No | *Sarax* sp. Bali | Indonesia, Bali |
| LP 1926 | COI-MT040900 28S-KY509385 18S-KY509335 16S-MF806081 12S-MF806124 | No | *Sarax brachydactylus* Simon, 1892 | Philippines |
| LP 5564 | COI-MT040913 28S-MF806091 18S - MF806137; 16S -KY509397; 12S - KY509347 | No | *Sarax bilua* Miranda, Prendini, Giupponi & Scharff, 2021 | Solomon Islands |
| LP 13118 | COI-MT040902 28S-KY509387 18S - KY509337; 16S - MF806083; 12S - MF806126 | No | *Sarax cochinensis* (Gravely, 1915) | Kerala - India |
| LP 12298 | COI-MT040899 28S-KY509384 18S-KY509334 16S-MF806080 12S-MF806123 | No | *Sarax bispinosus* (Nair, 1934) | India, Sri Lanka |
| MNHN-JAC60 | COI-MT040899 28S-KY509384 18S-KY509334 16S-MF806080 12S-MF806123 | No | *Catageus* sp.3 | Unknown |
| MR365 (unregistered) | UCE, this study | No | *Charon* sp. | Torres Strait: Dauan Island |
| MNHN-JAA95 | COI-JN018110 28S-JN018324 18S-JN018227 | No | *Sarax* sp. | Unknown |
| LP 11998 | COI-MT040935 28S-MF806109 18S-MF806160 16S-KY509420 12S-KY509370 | No | *Sarax tiomanensis* Miranda, Prendini, Giupponi & Scharff, 2021 | Tioman Island - Malaysia |
| LP 11997 | COI-MT040924 28S-KY509409 18S-KY509359 16S-MF806102 12S-MF806149 | No | *Sarax rimosus* (Simon, 1901) | Malaysia, Singaporae |
| LP 1927 | COI-MT040898 28S-KY509383 18S-KY509333 16S-MF806079 12S-MF806122 | No | *Sarax batuensis* Roewer, 1962 | Batu Caves |
| MNRJ | UCE, this study | No | *Charinus diamantinus* Miranda, Prendini, Giupponi & Scharff, 2021 | caves of Diamantina Plateau - Brazil |
| LP 10076 | COI-MT040903 28S-KY509388 18S-KY509338 16S-MF806084 12S-MF806127 | No | *Charinus gertschi* Goodnight & Goodnight, 1946 | Guyana, Surinam, Venezuela |
| LP 13448 | COI-MT040929 28S-KY509414 18S-KY509364 12S-MF806154 | No | *Charinus sillami* Réveillion & Maquart, 2015 | French Guyana |

| | | | | |
|---|---|---|---|---|
| LP 3831 | COI-MT040917 28S-KY509401 18S-KY509351 16S-MF806095 12S-MF806141 | No | *Charinus palikur* Miranda, Prendini, Giupponi & Scharff, 2021 | French Guiana, Roura |
| LP 6943 | COI-MT040897 28S-KY509382 18S-KY509332 16S-MF806078 12S-MF806121 | No | *Charinus africanus* (Hansen, 1921) | Equatorial Guinea, São Tomé and Principe |
| ZMH: A0000893 | COI-MH107031 | No | *Charinus kakum* Harms, 2018 | Kakum National Park - Ghana |
| LP 13396 | COI-MT040939 28S-KY509424 18S-KY509374 16S-MF806113 12S-MF806164 | No | *Charinus vulgaris* Miranda & Giupponi, 2011 | Porto Velho - Rondonia, Salvador - Bahia - Brazil |
| USNMENT 1458725 | UCE, this study | No | *Charinus* sp.2 | Brazil, Pernambuco, Recife |
| LP 10473 | COI-MT040906 28S-KY509390 18S-KY509340 12S-MF806129 | No | *Charinus dominicanus* Armas & Pérez, 2001 | Dominican Republic |
| LP 10170 | COI-MT040938 28S-KY509423 18S-KY509373 16S-MF806112 12S-MF806163 | No | *Charinus aguayoi* Moyá-Guzman, 2009 | Puerto Rico |
| | SRX10297721 | No | *Charinus acosta* (Quintero, 1983) | Cuba |
| LP 13402 | COI-MT040922 28S-KY509407 18S-KY509357 16S-MF806100 12S-MF806147 | No | *Charinus reddelli* Miranda, Giupponi & Wizen, 2016 | Footprint Cave and Waterfall Cave - Belize |
| UFPI 11323 | UCE, this study | No | *Charinus carvalhoi* Miranda, Prendini, Giupponi & Scharff, 2021 | Roraima - Brazil |
| MNRJ | UCE, this study | No | *Charinus magalhaesi* Miranda, Prendini, Giupponi & Scharff, 2021 | Amazonas, Manaus - Brazil |
| LP 6366 | COI-MT040920 28S-KY509405 18S-KY509355 16S-MF806099 12S-MF806145 | No | *Charinus pescotti* Dunn, 1949 | Australia, Queensland |
| ARALP000157/ LP 10276/ LP 5174 | H3-KY018133 COI-MT040916 28S-KY509400 18S-KY509350 16S-MF806093 12S-MF806139 | No | *Charinus neocaledonicus* Kraepelin, 1895 | New Caledonia |
| LP 5175 | 28S-KY509391 18S-KY509341 16S-MF806086 12S-MF806130 | No | *Charinus elegans* (Weygoldt, 2006) | New Caledonia |
| USNMENT 1458727 | UCE, this study | No | *Charinus una* Miranda, Prendini, Giupponi & Scharff, 2021 | Brazil, Bahia |
| LP 13400 | COI-MT040934 28S-KY509419 18S-KY509369 12S-MF806159 | No | *Charinus taboa* Vasconcelos, Giupponi & Ferreira, 2016 | Brazil, Minas Gerais |
| Charinus_6.3 Roc | COI-MK801767 16S-MK810724 | No | *Charinus rocamadre* Torres-Contreras, García & Armas, 2015 | Colombia, Sucre |

| | | | | |
|---|---|---|---|---|
| MNRJ 9366 | UCE, this study | No | *Charinus montanus* Weygoldt, 1972 | Brazil, Espírito Santo |
| AM7 | 28S-KP276344 18S-KP276424 | No | *Charinus asturius* Pinto-da-Rocha, Machado & Weygoldt, 2002 | Brazil, São Paulo |
| USNMENT01408166 | UCE, this study | No | *Charinus euclidesi* Miranda, Prendini, Giupponi & Scharff, 2021 | Brazil, Rio de Janeiro |
| MNRJ | UCE, this study | No | *Charinus caatingae* Vasconcelos & Ferreira, 2016 | Brazil, Bahia |
| LP 13398 | COI-MT040921 28S-KY509406 18S-KY509356 12S-MF806146 | No | *Charinus potiguar* Vasconcelos, Giupponi & Ferreira, 2013 | Brazil, Rio Grande do Norte |
| MNRJ | UCE, this study | No | *Charinus* cf *potiguar* | Brazil, Rio Grande do Norte |
| USNMENT01408154 | UCE, this study | No | *Charinus souzai* Miranda, Prendini, Giupponi & Scharff, 2021 | Brazil, Espírito Santo |
| USNMENT01408153 | UCE, this study | No | *Charinus sooretama* Miranda, Prendini, Giupponi & Scharff, 2021 | Brazil, Espírito Santo |
| USNMENT01580325 | UCE, this study | No | *Charinus* sp.1 | Brazil, Espírito Santo |
| MNRJ 9363 | UCE, this study | No | *Charinus brasilianus* Weygoldt, 1972 | Brazil, Espírito Santo |
| USNMENT01580322 | UCE, this study | No | *Charinus* sp.3 | Brazil, Espírito Santo |
| MNRJ 9362 | UCE, this study | No | *Charinus ruschii* Miranda, Milleri-Pinto, Gonçalves-Souza, Giupponi & Scharff, 2016 | Brazil, Espírito Santo |
| USNMENT01408205 | UCE, this study | No | *Charinus goitaca* Miranda, Prendini, Giupponi & Scharff, 2021 | Brazil, Rio de Janeiro |
| USNMENT01580327 | UCE, this study | No | *Charinus* sp. | Brazil, Espírito Santo |

**Supplementary Table S3.** Fossil calibrations and references for divergence time estimation.

| Node | Fossil | Taxonomy | Calibration | Node calibrated | Reference |
|---|---|---|---|---|---|
| **Root** | *Eramoscorpius brucensis* | Scorpiones (stem-group) | 435 Mya (lower) to 545 (upper) | Root | Waddington et al. 2015 |
| **Crown Scorpiones** | *Compsoscorpius buthiformis* | Scorpiones, crown Orthosterni | 314.6 (lower) | MRCA (Scorpiones) | Pointon et al. 2012 |
| **Crown Araneae** | *Palaeothele montceauensis* | Araneae, stem-Mesothelae | 299 (lower) | MRCA (Araneae) | Selden 1996 |
| **Crown Uropygi** | *Parageralinura naufraga* | Thelyphonida, stem-Uropygi | 319 (lower) | MRCA (Thelyphonida) | Brauckmann and Kock 1983 |
| **Crown Tetrapulmonata** | *Attercopus fimbriunguis* | Uraraneida | 374 (lower) | MRCA (Tetrapulmonata) | Selden et al. 2008 |
| **Crown Amblypygi** | *Weygoldtina angelica* | Amblypygi, stem-Palaeoamblypygi | 313.7 (lower) | MRCA (Amblypygi) | Pocock 1911 |
| **Crown Charontida** | *Kronocharon prendinii* | Amblypygi, Charontida | 105 (lower) | MRCA (Charontida) | Grimaldi and Engel 2014 |
| **Crown Phrynida** | *Britopygus weygoldti* | Amblypygi, Phrynida | 112 (lower) | MRCA (Phrynida) | Dunlop and Martill 2002 |

**Supplementary Table S4.** Concatenation of UCE and six-marker data set.

| Sanger names | UCE names |
|---|---|
| Acanthophrynus_coronatus | Acanthophrynus_coronatus |
| Catageus_sp1 | |
| Catageus_sp2 | |
| Catageus_sp3 | |
| Centruroides_sculpturatus_AR105 | Centruroides_sculpturatus_AR105 |
| Chaerilus_celebensis | Chaerilus_celebensis |
| Charinus_acosta | Charinus_acosta |
| Charinus_africanus | |
| Charinus_aguayoi | |
| Charinus_asturius | Charinus_asturius_GSM243 |
| Charinus_brasilianus_GSM159 | Charinus_brasilianus_GSM159 |
| Charinus_caatingae_GSM373 | caatingaeGSM373 |
| Charinus_carvalhoi | |
| Charinus_diamantinus3762 | charinus_diamantinusgsm376_2 |
| Charinus_dominicanus | |
| Charinus_elegans | |
| Charinus_euclidesi_GSM257 | Charinus_euclidesi_GSM257 |
| Charinus_gertschi | |
| Charinus_goitaca_GSM133 | Charinus_goitaca_GSM133 |
| Charinus_kakum | |
| Charinus_magalhaesi | Charinus_magalhaesi_GSM394 |
| Charinus_montanus_GSM123 | Charinus_montanus_GSM123 |
| Charinus_neocaledonicus | |
| Charinus_palikur | |
| Charinus_pescotti | |
| Charinus_potiguar_GSM371 | Charinus_potiguar_GSM371 |

| | |
|---|---|
| Charinus_reddelli | |
| Charinus_rocamadre | |
| Charinus_ruschii_GSM240 | Charinus_ruschii_GSM240 |
| Charinus_seychellarum | Charinus_seychellarum_GSM413 |
| Charinus_sillami | |
| Charinus_sooretama_GSM233 | Charinus_sooretama_GSM233 |
| Charinus_souzai_GSM231 | Charinus_souzai_GSM231 |
| Charinus_sp._1196 | Charinus_sp._1196 |
| Charinus_sp1_GSM173 | Charinus_sp1_GSM173 |
| Charinus_sp2_GSM386 | Charinus_sp2_GSM386 |
| Charinus_sp3_1181 | Charinus_sp3_1181 |
| Charinus_taboa | |
| Charinus_vulgaris | Charinus_vulgaris_GSM391 |
| Charon_CMB043 | Charon_sp._CMB043 |
| Damon_annulatipes | |
| Damon_diadema | |
| Damon_gracilis | |
| Damon_johnstonii | |
| Damon_medius | |
| Damon_sylviae | |
| Etienneus_africanus | |
| Euphrynichus_bacillifer | Euphrinychus_bacillifer_clean |
| Heterophrynus_alces | |
| Heterophrynus_batesii | |
| Heterophrynus_longicornis | |
| Heterophrynus_origami_CHNUFPI2141_CMB085 | Heterophrynus_origami_CHNUFPI2141_CMB085 |
| Heterophrynus_sp._CHNUFPI2271_CMB090 | Heterophrynus_sp._CHNUFPI2271_CMB090 |
| Heterophrynus_sp._CMB083 | Heterophrynus_sp._CMB083 |
| Liphistius_malayanus | Liphistius_yangae |
| Mastigoproctus_giganteus | Mastigoproctus_giganteus |
| Musicodamon_atlanteus | Musicodamon_atlanteus_GSM416-et-422 |
| Pandinus_imperator | Pandinus_imperator |
| Paracharon_sp._GSM424 | Paracharon_sp._GSM424 |
| Paraphrynus_aztecus | |

| | |
|---|---|
| Paraphrynus_baeops | |
| Paraphrynus_carolynae | |
| Paraphrynus_cubensis | |
| Paraphrynus_laevifrons | |
| Paraphrynus_mexicanus | |
| Paraphrynus_pococki | |
| Paraphrynus_raptator_GSM414 | Paraphrynus_raptator_GSM414 |
| Paraphrynus_robustus | |
| Paraphrynus_sp._AH5XA82 | Paraphrynus_sp._AH5XA82 |
| Paraphrynus_viridiceps | |
| Phrynichodamon_scullyi | |
| Phrynichus_orientalis | |
| Phrynichus_scaber | |
| Phrynus_longipes | |
| Phrynus_marginemaculatus | Phrynus_marginemaculatus |
| Phrynus_sp._AH5XA68 | Phrynus_sp._AH5XA68 |
| PhrynusAH5AX68 | |
| Protoschizomus_pachypalpus | |
| Sarax_batuensis | |
| Sarax_bilua | |
| Sarax_brachydactylus | |
| Sarax_buxtoni | |
| Sarax_cochinensis | |
| Sarax_cochinensis_bispinosus | |
| Sarax_ioanniticus | Sarax_ioanniticus |
| Sarax_israelensis | Sarax_israelensis |
| Sarax_rimosus | |
| Sarax_singaporae | |
| Sarax_sp_Bali | |
| Sarax_sp_JA2011 | |
| Sarax_tiomanensis | |
| Sarax_yayukae | |
| Schizomida_undet_SaoTome_Principe_GSM438 | Schizomida_undet_SaoTome_Principe_GSM438 |
| Schizomida_undet_Tanzania_GSM445 | Schizomida_undet_Tanzania_GSM445 |
| Stenochrus_portoricensis | Stenochrus_sp._IZ74289 |
| Stenochrus_sbordonii | |

| | |
|---|---|
| Thelyphonellus_amazonicus | |
| Trichodamon_princeps | Trichodamon_princeps_GSM423 |
| Typopeltis_crucifer | |
| unaGSM381 | Charinus_una_GSM381 |
| Weygoldtia_consonensis | |
| Weygoldtia_davidovi | |
| Xerophrynus_machadoi | |
| | Acanthoscurria_geniculata |
| | Heterophrynus_longicornis_CHNUFPI2142 |
| | Heterophrynus_sp._CMB086 |
