## Supplementary Fig. S1 for "The rediscovery of a relict unlocks the first global phylogeny of whip spiders (Amblypygi)"

**Supplementary Figure 1.** Phylogenetic relationships of Amblypygi reconstructed using UCE data at occupancies; a) 10%; b) 25%; c) 40%.

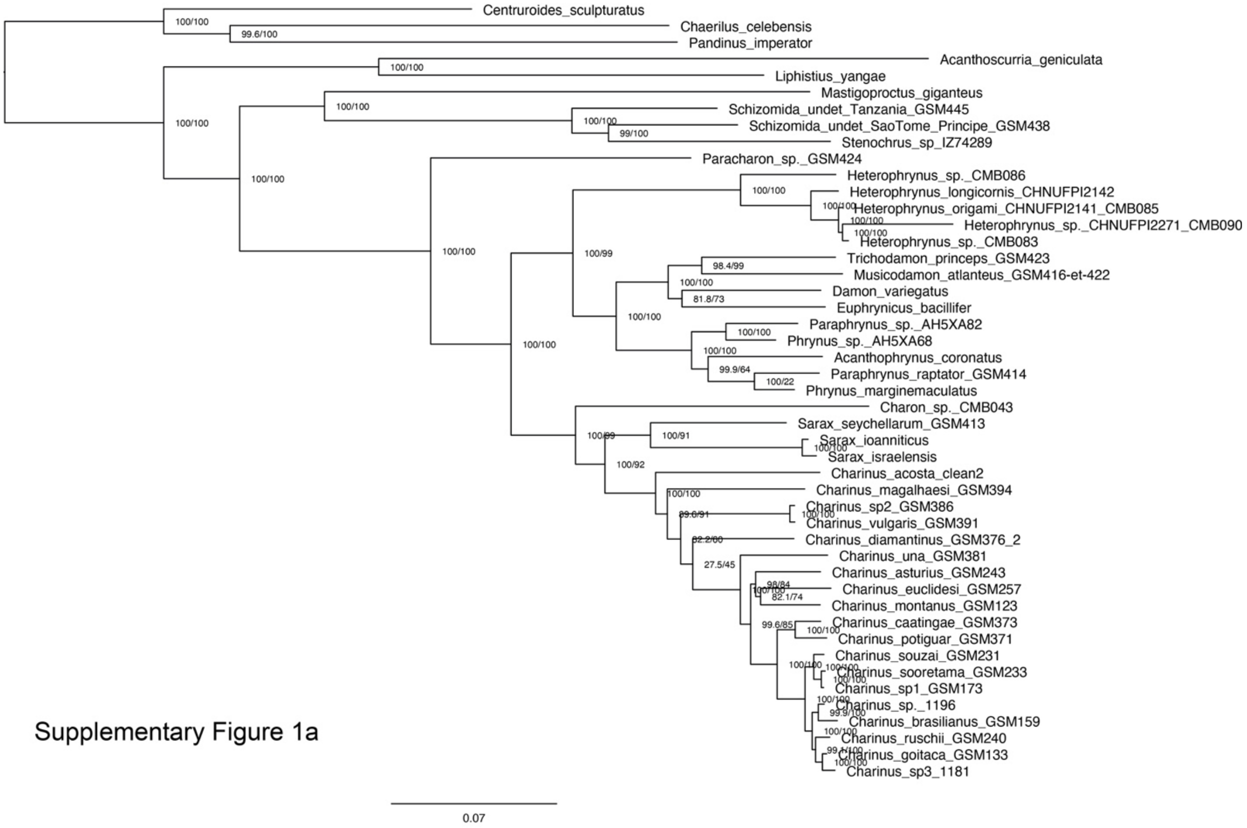


Supplementary Figure 1a

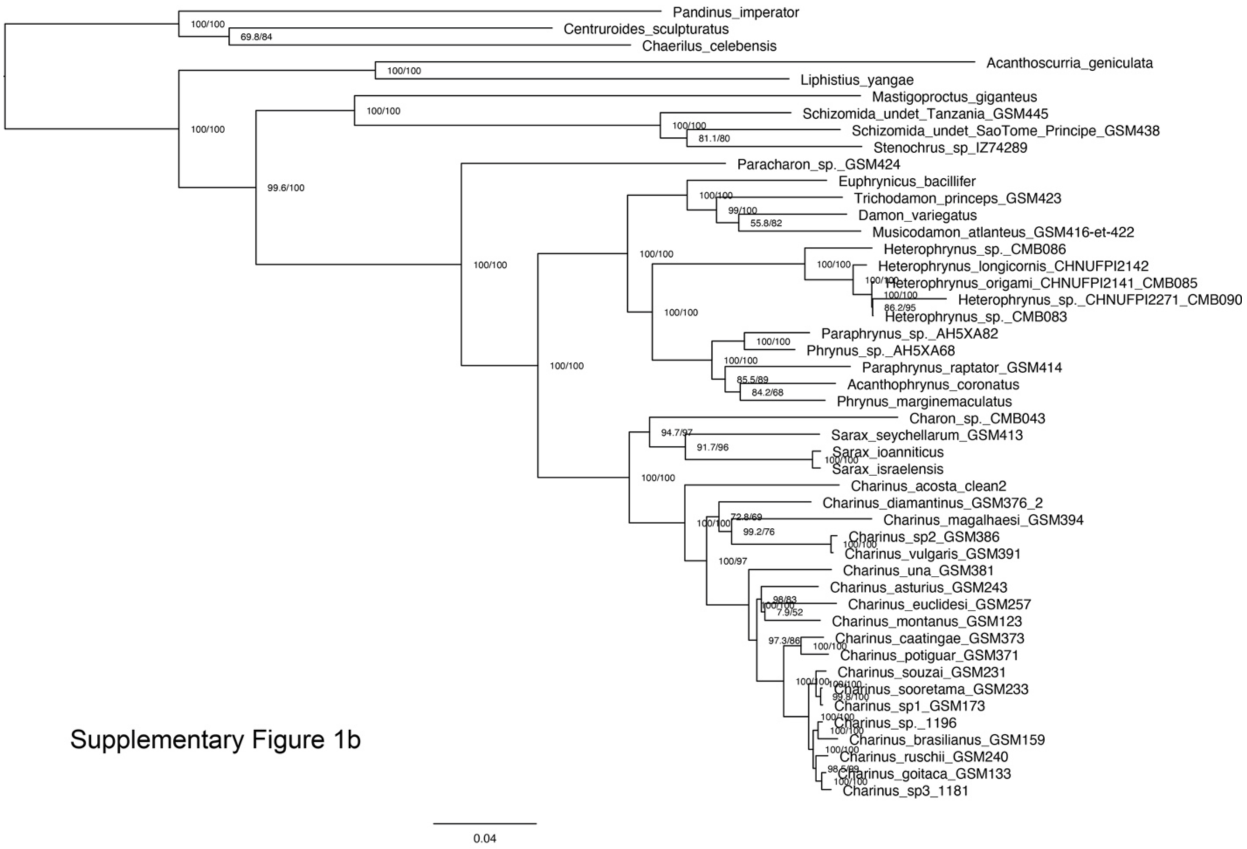


Supplementary Figure 1b

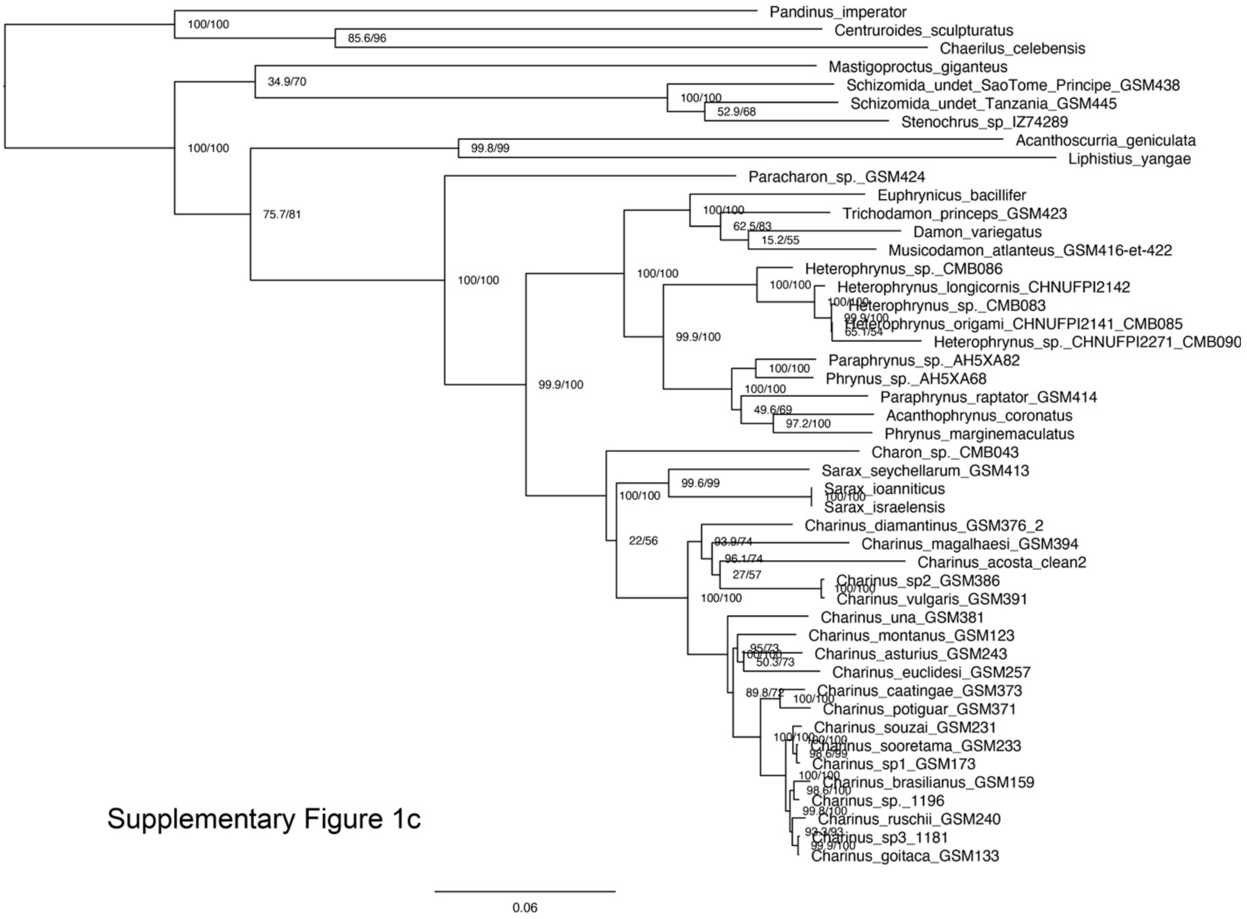

Supplementary Figure 1c
