## Supplementary Fig. S2 for "The rediscovery of a relict unlocks the first global phylogeny of whip spiders (Amblypygi)"

**Supplementary Figure 2.** Median node ages inferred by MCMCTree (top) and LSD2 (bottom), upon retention (purple) or exclusion (green) of Paracharontidae.

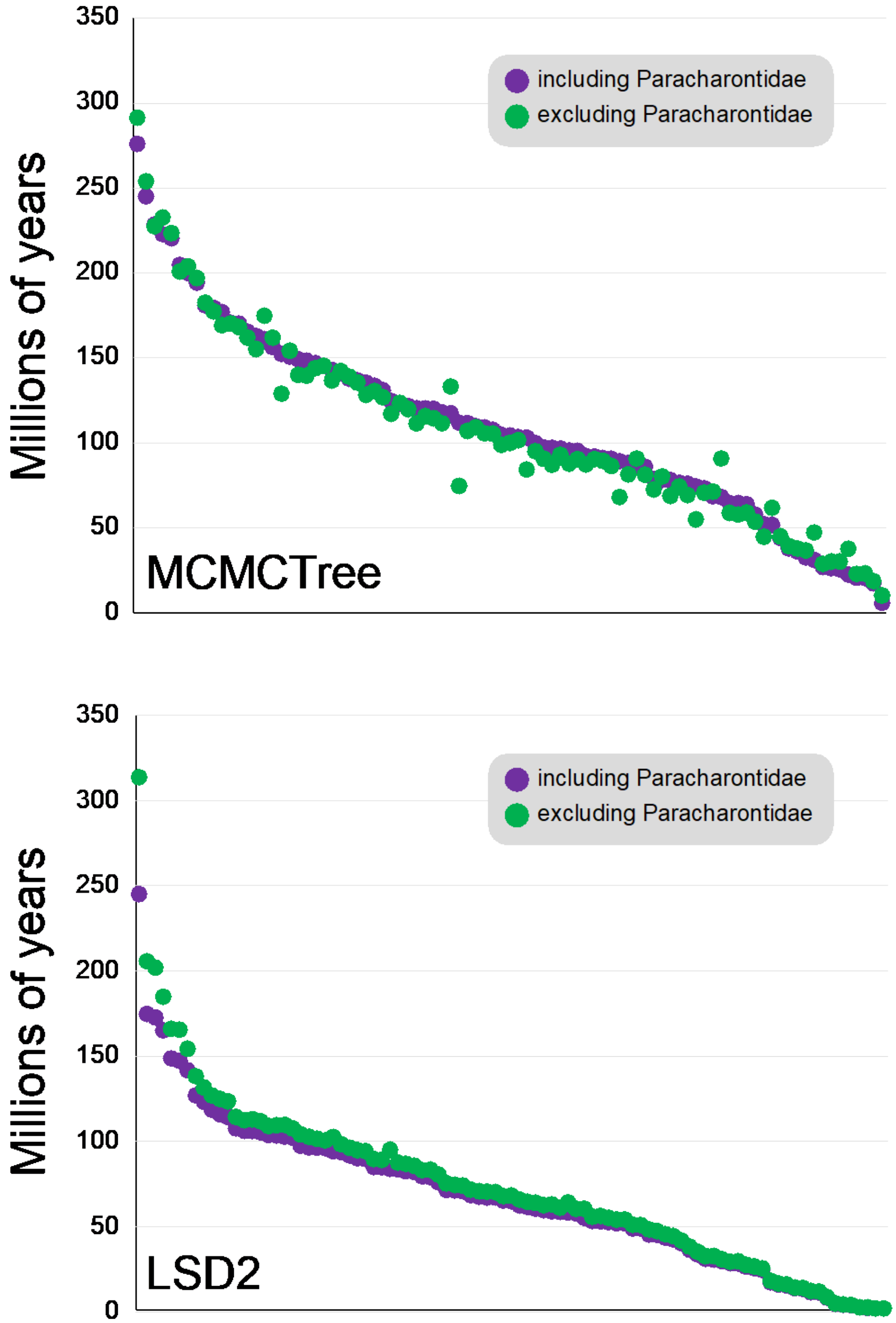
